# Structure and Interactions of the Endogenous Human Commander Complex

**DOI:** 10.1101/2023.04.03.535349

**Authors:** Saara Laulumaa, Esa-Pekka Kumpula, Juha T Huiskonen, Markku Varjosalo

## Abstract

The Commander complex, a 16-protein subunit assembly, plays multiple roles in various intracellular events, including regulation of cell homeostasis, cell cycle, and immune response. The complex is composed of COMMD1-10, CCDC22, CCDC93, DENND10, VPS26C, VPS29, and VPS35L. These proteins are expressed ubiquitously in the human body and have been linked to diseases including Wilson’s disease, atherosclerosis, and several cancers. Despite its importance, the structure and molecular functions of the Commander complex are poorly understood. Here, we report the structure and key interactions of the endogenous human Commander complex by cryogenic electron microscopy (cryo-EM) and mass spectrometry-based proteomics. Our results show that the complex is asymmetric, consisting of a stable core of a pseudo-symmetric ring of COMMD proteins 1–10 and a mobile effector consisting of DENND10 and the Retriever sub-complex, constituted by VPS35L, VPS29 and VPS26C. The two halves are scaffolded together by CCDC22 and CCDC93. This study directly confirms the cellular composition of Commander and identifies major interaction interfaces, defining the structure and interaction landscape of the complex. These findings offer new insights into its known roles in endosomal processes and intracellular transport, and uncovers a strong association with cilium assembly, and centrosome and centriole functions.

**Highlights:**

- High-resolution structure of the Commander complex determined by cryo-EM shows a rigid COMMD1–10 core decorated by mobile effectors DENND10 and the Retriever sub-complex.
- Comprehensive interactome analysis uncovers a plethora of novel interactions, implicating the Commander complex in previously unidentified processes, such as cilium assembly, cell cycle, and organelle biogenesis.
- Structural organization and identified interaction partners of the Commander complex provide a basis for further research into its molecular functions, related diseases, and potential therapeutic targets.

**eTOC blurb:** We provide a comprehensive structural analysis of the endogenous human Commander complex, revealing its functional organization and novel interactions involved in diverse cellular processes. These findings uncover new interaction surfaces and pave the way for further research into the molecular mechanisms of the Commander complex.

## Introduction

Cellular homeostasis maintenance in the trans-Golgi network is essential for key cellular functions, including cell growth, cell cycle, cell death, signal transduction pathways, and immune response ^1-3^. The recently discovered multiprotein Commander complex is a master regulator of cellular homeostasis within the trans-Golgi network ^4,5^. The existence of the Commander complex has been confirmed through protein-protein interaction (PPI) analysis ^2^ in addition to several contemporary interactome studies and phylogenetic profile analyses ^6-9^. Commander complex regulates cell signalling by affecting trafficking of transmembrane channel proteins and receptors to the cell surface and diverting them from lysosomal degradation. The complex has been linked to the regulation of several other cellular functions, including ion and lipid homeostasis ^10^ ^11^, embryogenesis ^4^, immune response ^9,12-16^, cell growth ^17^, and cell cycle ^16,18-20^.

The Commander complex consists of ten homologous Copper-metabolism Murr1 Domain proteins (COMMD1–10), coiled-coil domain-containing proteins CCDC22 and CCDC93, vacuolar protein sorting-associated proteins VPS26C, VPS35L, and VPS29 ^5,21^, and a DENN (differentially expressed in normal and neoplastic cells) domain-containing protein 10 (DENND10) ^8,22^. COMMD proteins 1–10 are sequence homologs of COMMD1, named after its first discovered function in regulating copper homeostasis, knockouts of which show defects in trafficking of copper transporters ATP7A and ATP7B ^23^. Later work has established that it controls sodium homeostasis through regulation of the human epithelial sodium channel (ENaC), plasma cholesterol levels through low-density lipoprotein receptor (LDLR), and chloride homeostasis through cystic fibrosis transmembrane conductance regulator (CFTR) ^24^, ^10,25^. Knockouts of individual COMMD proteins cause severe reduction in protein levels of all COMMD proteins, suggesting that they assemble together into a larger complex ^25^.

All ten COMMD genes in vertebrates are 90% conserved in mammals, while individual COMMD genes are found in genomes of lower metazoans like insects and worms. In lower vertebrates, knockdowns of COMMD proteins have been associated with developmental disorders ^4^. Similarly, in humans, the COMMD genes tolerate missense mutations poorly, as evidenced by the Genome Aggregation Database (gnomAD) ^26^. The existing few autosomal mutations in the COMMD genes have been linked to Wilson’s disease ^27^, Parkinson’s disease ^28^, atherosclerosis ^25^, and viral protein recycling ^21,29^. Additionally some somatic mutations are linked to cancer ^30-32^.

The COMMD proteins contain two domains: a C-terminal COMM domain unique to this family and a globular α-helical N-terminal domain (NTD), both individually solved by X-ray crystallography and nuclear magnetic resonance ^33-35^ ^35^. COMMD6, as an exception, contains a COMM domain but lacks the NTD. The C-terminal COMM domain forms an elongated structure that dimerizes via a left-handed handshake interaction ^34^. *In vitro*, COMMD proteins form elongated homo- and heterodimers ^34,36^. The CCDCs belong to a large family of coiled-coil domain containing proteins that have wide-ranging roles in the cell from ciliary motility and cell division to centrosome organisation ^37^. CCDCs often serve as scaffolds in larger protein complexes. CCDC22 in particular has been associated with X-linked intellectual disability / Ritscher-Schinzel syndrome / 3C syndrome ^14,38^. Both CCDC22 and CCDC93 contain an N-terminal microtubule binding NDC80 and NUF2 calponin-homology domain (NN-CH) and a C-terminal coiled-coil region. Knockdown of either CCDC causes defects in trafficking of transmembrane receptors such as α5β1-integrins ^21^.

DENND10 is a member of the DENND protein family of guanine nucleotide exchange factors (GEF) targeting Rabs. They constitute a ubiquitous, but relatively understudied family in metazoans with at least 18 members in 8 groups in humans ^39^. DENND10 is involved in late endosome homeostasis and exosome biogenesis, and has been proposed to interact with Rab27A or Rab27B ^40^.

The Commander complex is functionally linked to two other multiprotein complexes, the Retriever complex and the Retromer, which also play essential roles in endosomal sorting and cellular homeostasis. The Retriever is a recently discovered heterotrimeric complex composed of VPS29, VPS35L and VPS26C (or DSCR3) ^21^. It is thought to play a role similar to, but separate from, the homologous Retromer, which acts as a master controller of cargo sorting in eukarya. Retromer, consisting of VPS29, VPS35 and VPS26A/B associates with phosphatidylinositol-3-phosphate (PI(3)P) coated membranes on endosomes via protein coats formed by sorting nexins (SNX) SNX1/2 and SNX5/6 ^41,42^ and their phox homology domains (PH). Retromer forms a loosely symmetric coat composed of head-to-head retromer dimers that associate with the SNX coat via VPS26A/B and assist in formation of tubular transport intermediates ^43^. Retromer associates with auxiliary SNX proteins such as SNX3 or SNX27 that serve a cargo recognition function ^44^. SNX17 has been proposed as a Retriever-specific homolog of SNX27 in cargo recognition, but interactions between Retriever and other SNX proteins have not been demonstrated ^21^.

A central interacting partner for Retromer and Retriever is the heteropentameric Wiskott-Aldrich syndrome proteins and SCAR homologue (WASH) complex, an endosome-specific Arp2/3 complex activator that induces actin patch formation on endosomes ^21^ ^45,46^. These interactions are thought to be mediated by unstructured regions in WASH complex component WASHC2A either directly with VPS35 (or VPS35L in Retriever), or via SNX27 (or SNX17 in Retriever) ^21^.

Despite several studies focusing on individual Commander subunits, further systematic studies that assess the Commander complex as a whole are necessary to understand how the multitude of its different functions are regulated. Similarly, the atomic structure of the Commander complex remains elusive, hindering our understanding of its functions. Furthermore, in the absence of the structure, the exact mechanism behind the assembly of these proteins into a functional Commander complex is not yet fully understood.

In this study, we seek to address critical gaps in our understanding of the Commander complex. We report the structure of the endogenous human Commander complex determined by cryogenic electron microscopy (cryo-EM) and comprehensively map the molecular context and interactions of the Commander and individual complex interactions in human cells using affinity purification mass spectrometry (AP-MS) and proximity labelling (BioID). These findings provide a structure-based blueprint for understanding the functions of the Commander complex in endosomal sorting and other cellular processes. By elucidating the interactions between the complex components and other cellular constituents, this study sheds light on the Commander complex’s role in regulating cellular homeostasis, cell cycle, and immune response. In conclusion our study offers new insights into Commander’s functions and advances our understanding of its molecular mechanisms.

## Results

### Characterization of the composition and molecular context of the endogenous human Commander complex

To investigate the structure and molecular context of the endogenous human Commander complex, we employed a systematic affinity purification/mass spectrometry (AP-MS) and BioID workflow combined with cryo-EM structure determination (**Fig. 1A**). Isogenic Flp-In™ 293 T-REx cell lines were generated, with inducible expression of the individual 14 Commander complex components tagged with MAC-tags (StrepIII, HA, and BirA*) or StrepIII-tag, at either the N- or C-terminus (totaling 19 bait proteins), respectively (**Fig. 1B**). Single-step AP-MS and BioID were used to identify interacting proteins, with stringent statistical filtering applied to infer high-confidence interactions (HCIs).

**Figure 1.**
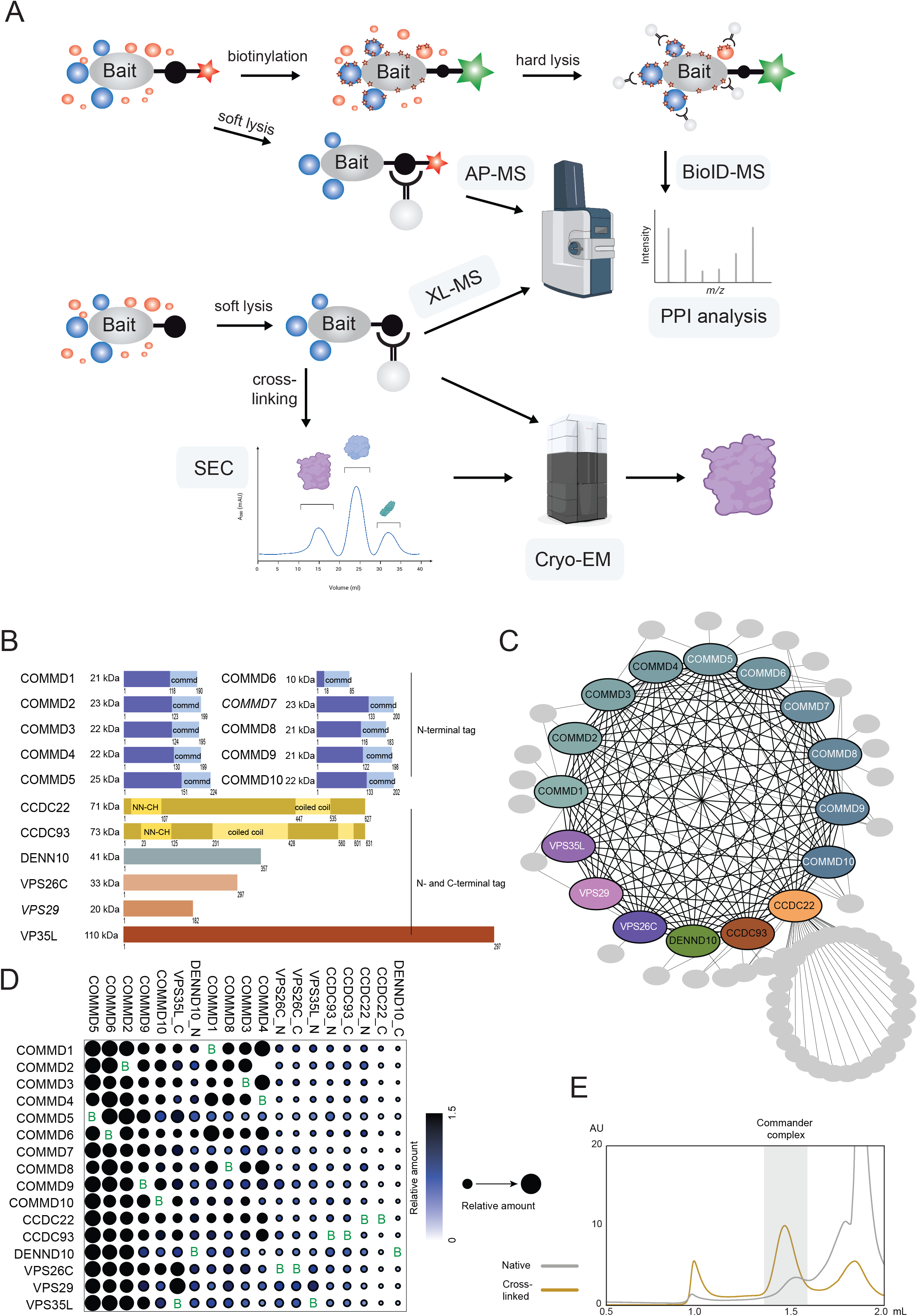
Purification and analysis of the endogenous Commander complex (**A**) Schematic of the study design utilising affinity purification mass spectrometry, proximity-dependent biotin identification, cross-linking mass spectrometry, size exclusion chromatography and cryogenic electron microscopy. (**B**) The known 16 members of the Commander complex proteins, their sizes (kDA) and known domain compositions. The 14 complex proteins used as baits in the studies are shown with normal typeface. (**C**) The AP-MS analysis identified high-confidence and stable Commander complex interactome. (**D**) Stoichiometry analysis of the Commander complex components identified with AP-MS. The colour of each circle represents the relative abundance of each prey (normalized to the mean abundance of the complex components for each bait), and the circle size indicates the relative abundance across all conditions. The relative abundance of each COMMD protein displays a stoichiometric ratio of <1 unit/complex. (**E**) Size exclusion chromatography of the purified Commander complex with or without crosslinking. The peak indicated in grey background was used for Cryo-EM analysis.

Using AP-MS, we identified 75 high-confidence interactors and 404 interactions for the 19 bait proteins (some tagged at the N- or C-terminus). The majority of interactions were with other Commander complex proteins, and we captured 93% of all the possible binary interactions within the complex—illustrating not only the ability of our method to retain intact protein complexes, but also the stability of the Commander complex (**Fig. 1C**). When we compared the 404 detected interactions with six interaction databases (BioGRID, IntAct, PINA2, String, bioplex, and human cell map), approximately 25% (115 interactions) were previously unreported, including uncharacterized interactions within the complex components and with other proteins (e.g. SNX9 and SNX27) and complexes (e.g. WASH and HAUS) (**Fig. S1**; **Table S1**). Interestingly many of the interactions with WASH proteins are detected with C-terminally tagged CCDC22 (**Fig. S1B**) The CDCC22 interaction abundances with the WASH proteins is lower than with Commander complex proteins or with PCM1 (Pericentriolar Material 1) and MTMR2 (Myotubularin-related protein 2), proteins which are required for WASH recruitment (**Fig. S1B**). The PCM1 is a centrosomal coiled coil protein, which appears to anchor the WASH complex at the centrosome, where it promotes the nucleation of branched actin networks (26655833). Whereas, the MTMR2 is recruited by the CCC complex to modulate the levels of phosphatidylinositol 3-phosphate (PI(3)P) on the membranes of endosomes which in turn recruits the WASH complex (31537807). Additionally, the other 30 interacting proteins appear specific for the CCDC22. Further Reactome pathway analysis of these proteins showed over 20-fold enrichment of R-HSA-5617833∼Cilium Assembly pathway, with 9 (30% of the total pathway components) proteins (OFD1, PCM1, NDE1, HAUS4, EXOC4, EXOC6, EXOC5, HAUS5, HAUS1) detected (**Table S2**).

In addition to identifying high-confidence interactions, we quantified the relative amount of the proteins within the complex to obtain the stoichiometries for the complex components. Normalized relative copy amounts of each Commander component purified with the 19 bait proteins are shown in **Fig. 1D**, and the baits are hierarchically clustered based on their similarity in interaction abundances. The COMMD proteins are present in close to a one molecule stoichiometries in the complex. This suggests that the COMMD proteins form a stable complex, predominantly containing all of the ten proteins. Interestingly, the COMMD proteins do still display two slightly separating clusters, with COMMD5, 6, 2, 9, and 10 in one and COMMD1, 8, 3, and 4 in the other, possibly corresponding to two regions within the structure. Also the VPS35L and DENND10 display to be predominantly associated with the complex, whereas the association of the CCDCs appears less frequent. The interaction within the Commander complex might be PTM regulated, therefore we searched for possible post transcriptional modifications (PTMs) from the AP-MS data. This identified several phosphosites, in the six of the complex components, and additionally histidine methylation in five components (**Table S2; Fig. S5A**).

### Structure of the endogenous human Commander complex

We purified the native Commander complex for structural analysis by Strep-Tactin affinity purification using tagged COMMD9 as a bait. A single step affinity purification yielded the complex in high purity, directly applicable for cryo-EM single particle analysis (SPA). We collected an initial cryo-EM dataset for the native Commander from affinity purified material. Based on the data, we reconstructed a cryo-EM density map at a nominal resolution of 3.4 Å, corresponding to the COMMD ring as well as parts of CCDC22 and CCDC93 (**Fig. 2A**, **Fig. S2A,C, Fig. S5B**). Presence of additional features in the 2D class averages (**Fig. S2A**) suggested that additional structures remained unresolved in the 3D density map, likely due to conformational heterogeneity. In order to stabilise the complex, we treated affinity purified Commander with bis(sulfosuccinimidyl)suberate (BS3) cross-linker in mild conditions. This treatment allowed purification of the complex by size exclusion chromatography (**Fig. 1E**). We obtained a peak corresponding to the expected size of the complex (560 kDa) which was used both for cross-linking mass spectrometry (XL-MS) and cryo-EM.

**Figure 2.**
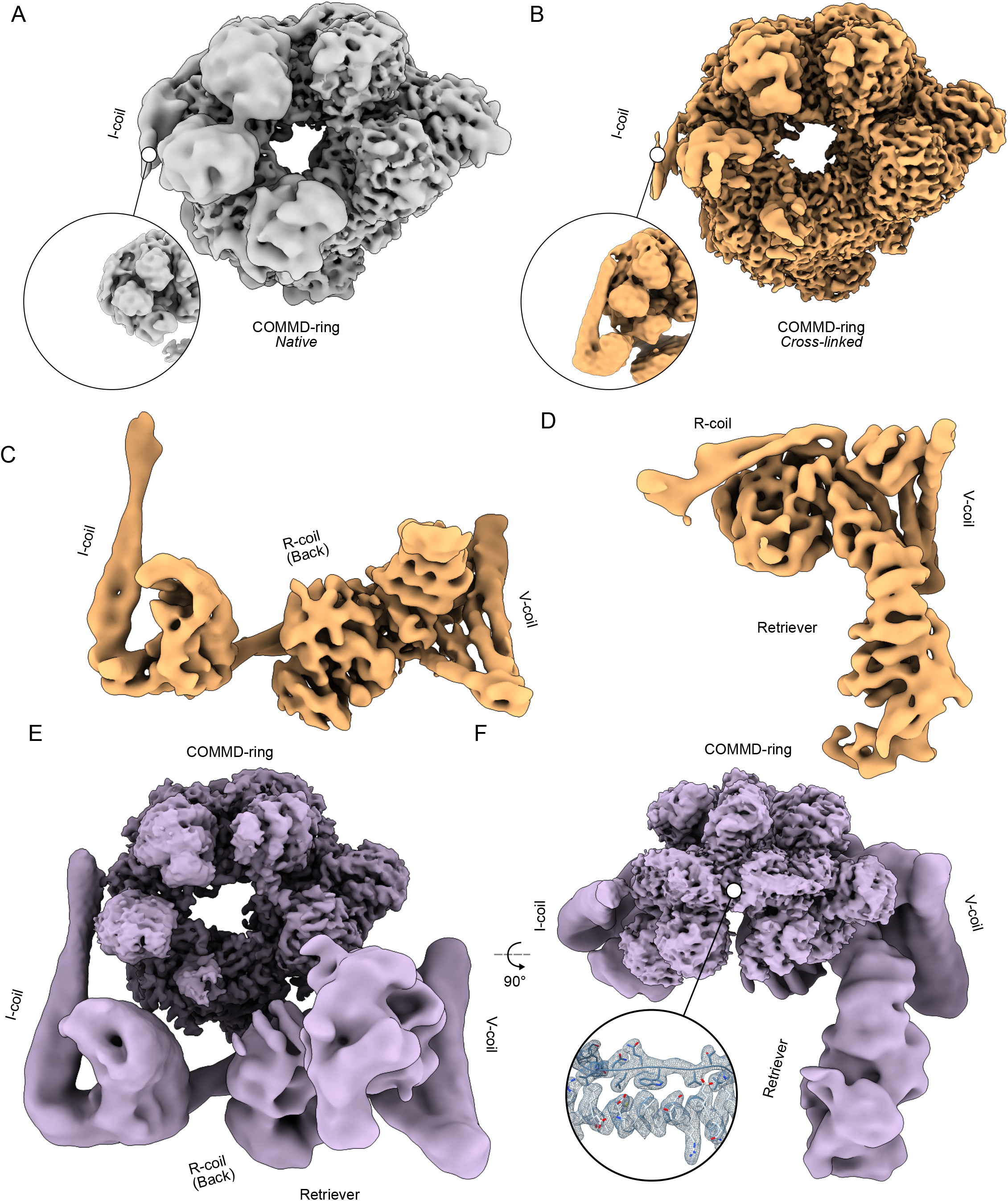
Cryo-EM maps of the Commander complex. (**A–B**) Cryo-EM maps of the Commander complex COMMD-ring from native (**A**, in gray) and from cross-linked (**B**, in gold) samples. Insets show the I-coil region from both maps at lower isosurface threshold. (**C**) Focused cryo-EM density map with DENND10 and CCDC22/93 coiled coils (focused map 1). (**D**) Focused cryo-EM map with Retriever subregion (focused map 2). (**E**) A composite map of the complete Commander complex. (**F**) Same as in E, rotated 90° around the x-axis as indicated. Inset: structural detail of the map from within the COMMD ring.

We determined a cryo-EM density map of the cross-linked Commander complex at a nominal resolution of 2.9 Å (**Fig. 2B, Fig. S2D-E, Fig. S5C**). The features of this map, such as an extended density on the side of the COMMD ring corresponding to the I-coil, were improved when compared to the map determined from the native dataset (**Fig. 2A-B**). Furthermore, the map determined from the cross-linked dataset indicated that the structures below the COMMD ring showed less conformational heterogeneity than in the native map. The remaining heterogeneity still prohibited fitting of atomic models into this density, and therefore we proceeded to do focused classification and refinement using partially signal subtracted particles. First, we subtracted the COMMD ring region of the map and through iterative rounds of 3D classification and local refinement reconstructed a density map corresponding to the I, R and V-coil regions, DENND10 density as well as a part of the Retriever density at a nominal resolution of 6.5 Å (**Fig. 2C, Fig. S2F Fig. S5D**). Second, we subtracted the I-coil, half of R-coil and DENND10 regions from the density, and focused on the V-coil and Retriever regions. After extensive classification and local refinement, we obtained a density map that contained the complete Retriever subcomplex as well as the V-coil region at a nominal resolution of 7.5 Å (**Fig. 2D, Fig. S2G, Fig. S5E**). We further combined these three maps into a composite map of the entire complex (**Fig. 2E,F**).

### The COMMD ring is highly interconnected and stable

AP-MS indicated that the Commander complex contains a single copy of each subunit (**Fig. 1D**). Based on this information, we used AlphaFold2 (AF2 ^47-49^) using a single copy of each sequence to generate an initial atomic model for the entire complex and different sub-complexes. First, we predicted the structure of the heterodecameric COMMD ring without the N-terminal domains (NTDs), as it had the highest resolution that would allow identification of individual COMMD components in the density. AF2 predictions converged on a ring-like structure with varying orders of COMMD subunits around the ring. Based on the fact that COMMD6 does not have an N-terminal domain (NTD), we anchored the models to the cryo-EM density and determined the correct order of COMMD subunits in the ring by density fit. We used the best fitting model within the COMMD domains as a starting point and refined the model into density using a combination of ISOLDE ^50^ and PHENIX ^51^ (**Fig. 3A**). To validate our assignment of different COMMD subunits, we tested placements of all COMMD proteins against all possible sites in the density. Ranking of these placements by model-to-map cross-correlation after refinement showed that our initial assignment was ranked the highest (**Fig. 3B-D**).

**Figure 3.**
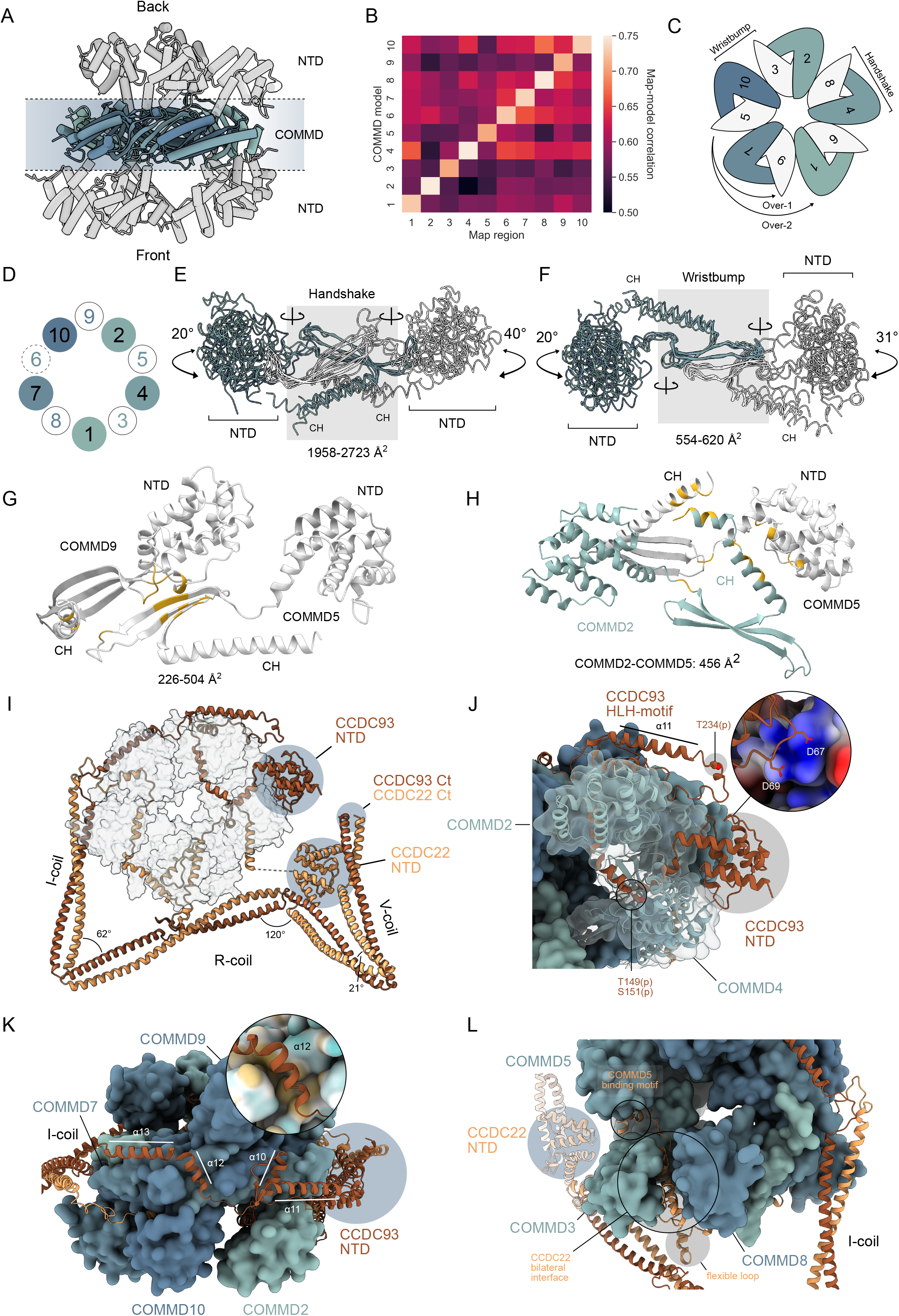
Structure of the COMMD ring. (**A**) The structure of the COMMD ring is shown as a ribbon representation from the side. COMMD domains and NTDs are indicated. (**B**) All vs. all heatmap depicting model-to-map cross correlation coefficients for COMMD domains placed in each COMMD map region after real space refinement in PHENIX. (**C**) Schematic diagram of COMMD ring subunit organization. COMMDs with NTDs towards the viewer are indicated in dark blue-green shades, while COMMDs with NTDs away from the viewer are indicated in white. Handshake, wristbump, over-1 and over-2 interactions are indicated with symbols. (**D**) Schematic representation of the NTD organization within the COMMD ring. Numbers indicate COMMD proteins. Coloring as in **C**. COMMD6 is indicated in dashed lines as it lacks an NTD. (**E**) Superposed COMMD chains are shown to view the handshake interaction between two chains (grey area) with range of interaction surface areas for each handshaking COMMD pair. Angles of NTD rotation along the indicated hinge axes are shown, indicating a larger variation of orientations on the back side of the COMMD ring. Color scheme as in **C**. (**F**) Similar to **E**, showing the wristbump interaction. (**G**) Model of over-1 interaction between COMMDs 2 and 5. Interacting residues indicated in gold, with buried surface area ranges indicated. (**H**) Model of over-2 interaction between COMMDs 5 and 2, color scheme as in **G**. (**I**) Overview of CCDCs intertwined within the COMMD proteins. (**J-L**) Major CCDC interaction sites with COMMD chains. (**J**) CCDC93 NN-CH domain binds to the side of the NTD of COMMD4 and encircles the NTD of COMMD2 in a headlock with HLH-motif contacting the NN-CH domain. Experimentally identified phosphorylation sites are indicated. (**K**) Top view of the COMMD ring, showing the arrangement of CCDC93 α10-α13. Inset: hydrophobic binding pocket of α12 on COMMD7. Coloring indicates hydrophobic (yellow) and hydrophilic (cyan) surfaces. (**L**) CCDC22 forms a bilateral interaction interface between COMMD3 and COMMD8. The loop protruding out of the cryo-EM density map between α13 and α14 of CCDC22 as well as the NN-CH domain of CCDC22 are indicated.

The COMMD ring exhibits pseudo-D5 symmetry, and is characterized by four types of interactions around the ring. We denote these interactions in order of descending buried interface area as handshake, wristbump, over-1, and over-2 interactions (**Fig. 3D-H, Fig. S3C-D**). Handshake (HS) interactions have been described in the literature ^34^, and constitute a large buried surface area (1958-2723 Å^2^) between two COMMD domains (**Fig. 3E**). This surface is mediated by the C-terminal helix (CH) and the three-stranded β-sheet of the COMMD fold clasping the opposing COMMD domain in a pseudo-C2-symmetric interaction. HS interactions are mainly mediated by sidechains. The wristbump (WB) interaction, in contrast, is defined by a much smaller interface (554-620 Å^2^), and consists of a highly curved intermolecular β-sheet, formed between the β1 of each COMMD. WBs are mainly mediated by backbone interactions that span from the absolutely conserved Trp on one COMMD to the respective Trp on the other COMMD (**Fig. S3B**). Both HS and WB interactions take place between COMMD subunits with NTDs pointing in opposite directions relative to the COMMD ring (**Fig. 3C**). The remaining two interfaces, over-1 and over-2, are smaller (<500 Å^2^). Over-1 interactions occur between COMMDs with NTDs on the same side of the ring, and localize on the linker between the COMMD domain and the NTD, and the β2–β3 loop contacting the β-sheet of the opposing COMMD subunit, as well as the CH contacting the β1–β2 loop (**Fig. 3G**). Only one significant over-2 interface exists, between COMMD2 and COMMD5 (456 Å^2^), which is due to the kink in the C-terminal α-helix of COMMD2. This in turn allows interactions between the CHs in addition to the interactions between CH of COMMD2 and the NTD of COMMD5 (**Fig. 3H**). These interactions together constitute a tightly bound core structure with several complementary interaction surfaces from one COMMD protein to up to 5 neighbouring COMMD proteins.

The NTDs of COMMDs 1, 7, 9 and 10 are less ordered than the rest. This is likely due to the flexibility of the linker between the domains. In addition, the NTDs of these COMMDs have the least stabilizing interactions with the CCDCs, with COMMD1 and COMMD9 having none, and COMMD7 and COMMD10 contacting only short helical segments of CCDC22 situated between flexible regions (**Fig. S3E**).

### COMMD ring is stabilized by interwoven CCDC93 and CCDC22 proteins

We next built models into the remaining areas of the cryo-EM density at 2.9-Å resolution. The predicted N-terminal NN-CH domain of CCDC93 fits well into the density on the side of the COMMD ring between NTDs of COMMD4 and COMMD5 and was used as a starting point to build the rest of the CCDC93 model (**Fig. 3I**). Through manual building and placement of high confidence prediction segments from AF2, we traced the CCDC93 model to the I-coil region of the map (residues 1–313; **Fig 3I**). We then used the same approach to build the middle region of CCDC22 (residues 109–322, **Fig 3I**). Finally, we validated the manually built parts of our model by comparing it to a model predicted by AF2 for the top part of the complex (**Fig. S3A**).

The COMMD ring can further be divided into two halves based on the positions of the CCDCs. Both CCDCs are heavily intertwined within the COMMD ring, but can be seen as taking different routes around the ring, with CCDC93 going anti-clockwise (4 and 2 in the front; 5, 9 and 6 in the back) and CCDC22 clockwise (1, 7 and 10 in the front; 5, 3 and 8 in the back; **Fig. 3D,I**). These roughly determine the top (CCDC93) and bottom (CCDC22) halves of the COMMD ring. Both CCDCs end up in the I-coil region where they form a heterodimeric coiled-coil that extends into the bottom half of the structure.

CCDC93 interacts with the NTDs of COMMD2 and COMMD4 (**Fig. 3I-J**). The NN-CH domain binds on the side of COMMD4 NTD, while α7 of CCDC93 binds an interface formed by α2–α3 loop, α4 and α7 of COMMD4. Many NTDs of COMMDs (2, 3, 4, 5, 7 and 10) bind a segment of the CCDCs via this interface, which we call the peptide binding (PB) site (**Fig. S3E**). The α3–α4 loop of CCDC93 contacts the extended CH of COMMD2 via electrostatics mediated by Asp67 and Asp69 (**Fig. 3J**). CCDC93 proceeds to circle around the NTD of COMMD2 in a headlock, binding the PB-site via α9, and forming a long helix-loop-helix motif (HLH; α10 and α11) that reaches all the way around to the NN-CH domain (**Fig. 3J**). The flexible loop region contains a phosphorylation site on Thr234 (**Fig. 3J, Table S2**). The protein chain then returns towards the I-coil region of the COMMD ring. The long loop in the HLH motif is flexible and therefore not visible in our cryo-EM map, but density features support our assignment of the two helices. The final helices (α12 and α13) predominantly bind the COMMD domain of COMMD7, with α12 binding in a tight hydrophobic pocket, before reaching the I-coil region (**Fig. 3K**).

CCDC22 occupies the lower half of the COMMD ring. Its N-terminal NN-CH domain is not visible in the consensus cryo-EM map, but can readily be identified in the focused maps (**Fig. 2C-D**). The first helix of CCDC22 in the consensus map is the α8, which binds the COMMD domains of COMMD2, COMMD3 and COMMD8 at the N-terminal end of its CH as well as parts of the β-sheet (**Fig. S3C**). It then binds the PB-site on COMMD5 via a short α-helix containing motif (**Fig. 3L, Fig. S3E**), before forming a bilateral binding interface between NTDs of COMMD3 and COMMD8 (**Fig. 3L**). At this point, a loop extends out of the COMMD ring and is not visible in the density map, indicating flexibility (**Fig. 3L**). Finally, α14 of CCDC22 binds COMMD domains of COMMD1, COMMD6 and COMMD8 in a similar topology to α8, before reaching the PB-sites on COMMD7 and COMMD10, and meeting CCDC93 at the I-coil region (**Fig. 3I, Fig. S3D**).

### CCDC22 and CCDC93 form a heterodimeric coiled coil scaffold that binds DENND10 and the Retriever subcomplex

Next, we built atomic models into the cryo-EM density of the bottom half of the Commander complex. The bottom half exhibits much more flexibility than the top half, as evidenced by the lower overall resolution of the focused maps (**Fig. 2C-D**). AF2 predictions, although getting the overall arrangement mostly correct, failed to reproduce a single model that would directly fit the density. Therefore, we generated the initial model in several parts and fitted them individually in the cryo-EM density before merging them to generate a complete model of this region (**Fig. S4A-C**).

The CCDCs form a long heterodimeric coiled coil with three flexible corners and three major coil regions that we call I-coil, R-coil and V-coil (**Fig. 4A**). Two high-confidence XL-MS crosslinks between Lys373 of CCDC22 and Lys352 of CCDC93 (I-coil) as well as Lys598 of CCDC22 and Lys601 of CCDC93 (V-coil) support the atomic model (**Fig. S4D, Table S2**). The I-coil begins close to the COMMD domains of COMMD1 and COMMD6. This assignment was further validated by XL-MS. Supporting our assignment, one crosslink was identified between Lys318 of CCDC93 and Lys114 of COMMD7 (**Fig. S4D, Table S2**). This crosslink is also likely partially responsible for the reduction in conformational heterogeneity upon crosslinking. Furthermore, another crosslink between the NTDs of COMMD2 (Lys42) and COMMD10 (Lys56) supports our assignment of the COMMD ring chains (**Fig. S4D, Table S2**).

**Figure 4.**
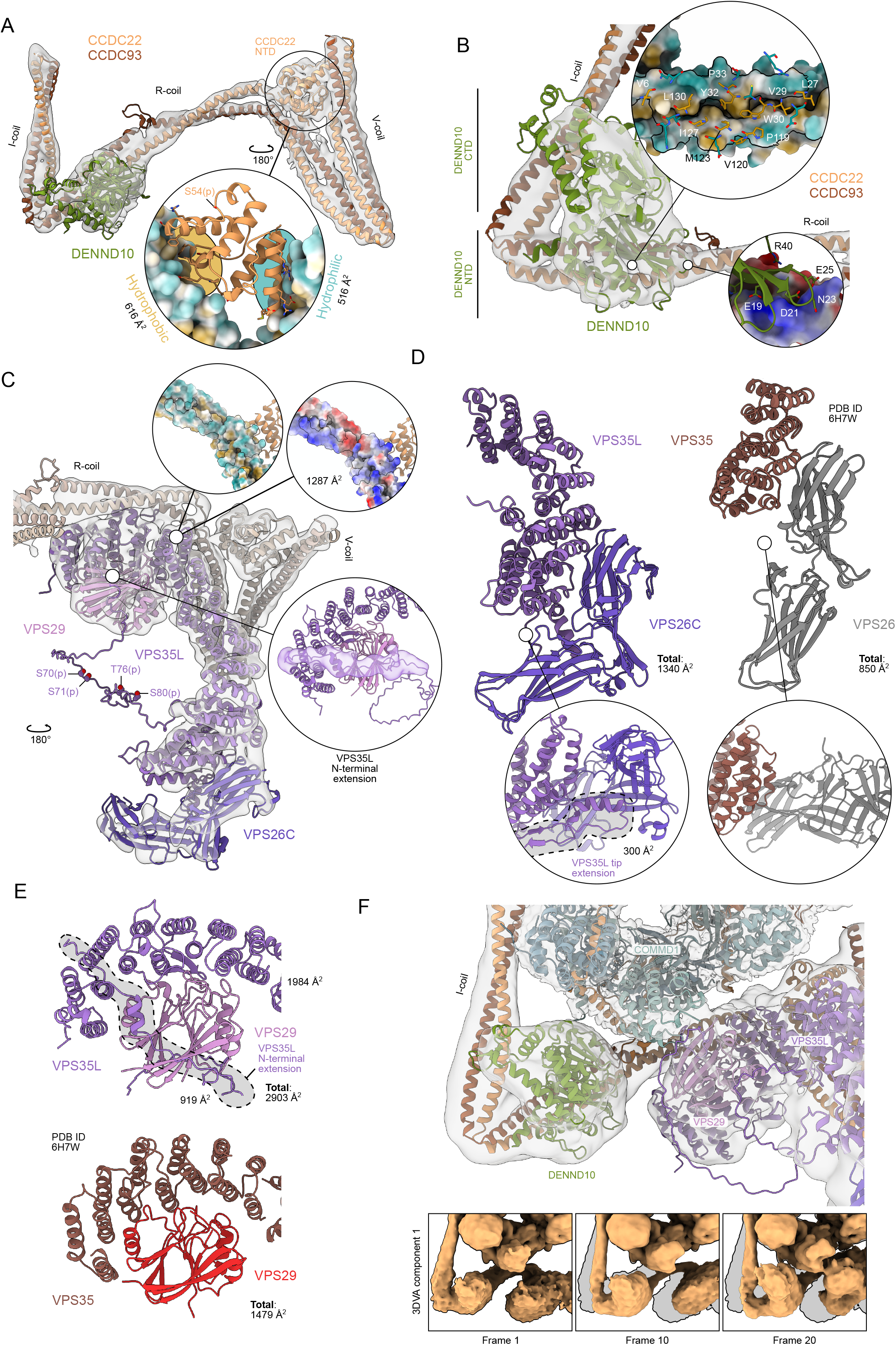
Structures and interactions of the effector subunits. (**A**) CCDC22/93 coiled-coil regions, NTD of CCDC22, and DENND10 in density. Inset: the binding site of CCDC22 NTD between the C-terminal V-coil characterised by hydrophobic and hydrophilic interfaces. Ser54 phosphorylation site identified in this study is indicated. (**B**) DENND10 interface with R-coil of CCDC22/93. The major interface consists of a hydrophobic core along the groove of the R-coil and a charged patch on the C-terminal side of R-coil. (**C**) The structure of the Retriever subcomplex in the context of Commander. Four experimentally identified phosphorylation sites (Ser70, Ser71, Thr76, and Ser80) are indicated in the disordered intervening N-terminal region of the model. Insets: Retriever interfaces with CCDC22/93 through the C-terminal part of VPS35L, mainly via charged interactions. The N-terminal extension of VPS35L is shown in density where contributions from VPS29 and VPS35L C-terminal parts have been masked and low-pass filtered. (**D**) Interfaces between the mobile NTD of COMMD1 with both DENND10 and VPS29/VPS35L. Inset: First, middle and last frames of component 1 from 3D variability analysis, reconstructed from particles using the intermediates job type. (**E**) Interaction between VPS35L and VPS29 (top) compared to corresponding interaction in fungal retromer arch (PDB 6H7W). The N-terminal extension of VPS35L accounts for nearly 33% of the buried interface. (**F**) Interaction between VPS35L and VPS26C compared to the fungal VPS35-VPS26 interface. Insets: VPS35L contains an extended helix that provides an expanded binding interface to VPS26C. Sidechains in (**A-B**) are included only for visualization purposes.

The NN-CH domain of CCDC22 is wedged between the C-terminal V-coil of CCDC93/22. The C-terminal half of the V-coil provides a hydrophobic binding interface, while the N-terminal half provides a hydrophilic one. In the complete model, the Cα–Cα distance between the last residue of the CCDC22 NTD and the first residue of α8 is 23 Å, a distance that accommodates the 12 residues in the linker between them. This is the only covalent link on this side of the complex between the top and bottom halves, which highlights the importance of the binding interface between the CCDC22 NTD and the V-coil in terms of flexibility of the bottom half. The phosphorylation site detected at Ser54 is unlikely to regulate this binding as it is located away from these surfaces, and is instead likely regulating external interactions (**Fig. 4A, Fig. S5A**).

The I-coil and R-coil form a 62° angle between them. The N-terminal side of R-coil harbors the binding site for DENND10 (**Fig. 4B**). The binding site is composed of two interaction surfaces, with a hydrophobic groove between the CCDC93/22 coil and the corresponding hydrophobic patch on the N-terminal domain (or lobe) of DENND10 comprising the main site, and one mediated by electrostatic interactions via Glu19, Asp21, Glu25 and Arg40 of DENND10 (**Fig. 4B**). Comparison to the structure of DENND1B in complex with Rab35 reveals that the putative Rab binding site of DENND10 is obstructed by the I-coil, indicating that DENND10 may be inactive in this conformation (**Fig. S4E**). However, the inherent flexibility of this region (**Fig. 4F**) may enable binding of Rabs in another conformation, or alternatively DENND10 may bind Rabs in an unconventional manner.

The AF2 model for Retriever is similar to that of Retromer, containing the distinctive α-solenoid fold (**Fig. S4C**). In the model of Retriever in the context of the Commander complex (**Fig. 4C**), the α-solenoid is more compacted and bent at the VPS26-binding end than in Retromer (**Fig. S4E**), shown by the decrease in distance of centres-of-mass of the ends (109 Å in Retromer, 91 Å in Retriever). In Retromer, the N-terminus is located at the VPS26-binding end, while the C-terminus is in the VPS29 binding end. In contrast to Retromer, Retriever has a long N-terminal extension of 180 residues. At the VPS26C binding end, the N-terminal extension contributes two additional helices (α2–α3). The tip extension (containing α3) provides a putative additional binding interface with VPS26C (**Fig. 4D**). We would like to note that relative to the entire map, the cryo-EM density in this area of the map is the weakest, suggesting conformational heterogeneity/structural flexibility, and the AF2 prediction confidence is the lowest. In the AF2 prediction, the N-terminal extension continues towards the VPS29-binding end of VPS35L but is unstructured in the intervening region. We find several phosphorylation sites in this region in our AP-MS data, indicating that this site could be a hotspot for phosphoregulation (**Fig. 4C, Fig. S5A**).

The very N-terminal part of the extension, the N-terminal tail (NTT), is predicted to contact VPS29 and reach all the way to the C-terminal part of VPS35L. This is supported by the density in our reconstruction, and we have therefore included the extension and the intervening unstructured region in our model (**Fig. 4C**). The N-terminal extension increases the binding interface between VPS35L and VPS29 significantly from 1984 Å^2^ to 2903 Å^2^. Compared to Retromer, this binding interface is nearly twice as large, suggesting tighter binding in Retriever (**Fig. 4E**). Retriever itself binds CCDCs at the hinge between R-coil and V-coil that forms a 120° angle, forming a large surface of mostly charged residues (**Fig. 4C**).

Finally, we computationally analysed structural flexibility within the complex. We found that despite seemingly not interacting with DENND10 or VPS29/VPS35L-NTT in the consensus cryo-EM map, COMMD1 NTD displays at least two alternative conformations where these interactions are imminent (**Fig. 4F**). This could indicate that the COMMD1 NTD has an important role in mediating interactions between Retriever and DENND10. The flexibility of the bottom half is facilitated by two mainchain contact sites at each end, namely via the I-coil and the linker between the NTD of CCDC22 and its top half portion. Other flexible joints are also present, including the tethers between I and R-coils as well as the R and V-coils. However, these are likely stabilised by binding of DENND10 and Retriever, respectively.

### Characterization of the cellular and molecular context of the Commander complex

To provide further insight into the molecular context, cellular interactions, and functions of the Commander complex, we focused on the HCI identified through BioID-MS analysis (see **Fig. 1; Fig. S6A**). In total, the BioID analysis resulted in the identification of 148 interacting proteins corresponding to 499 HCIs for the 14 Commander complex components used as baits. The ‘MS microscopy’ analysis showed a predominant localization for the complex proteins to Golgi, which is in agreement with the previous knowledge. Several components also displayed clear microtubule (COMMD8, COMMD4, COMMD3, COMMD2, COMMD1) and centrosomal (CCDC93 and CCDC22) localization (**Fig. S6B**). We compared our HCIs to previously reported interactions in six public interactome databases and observed that the majority of the interactions we identified were previously unreported (**Fig. 5A**, **Supplementary Data Table S1**). Our approach, therefore, provides a comprehensive view of the Commander complex interactions and sheds new light on its cellular role.

**Figure 5.**
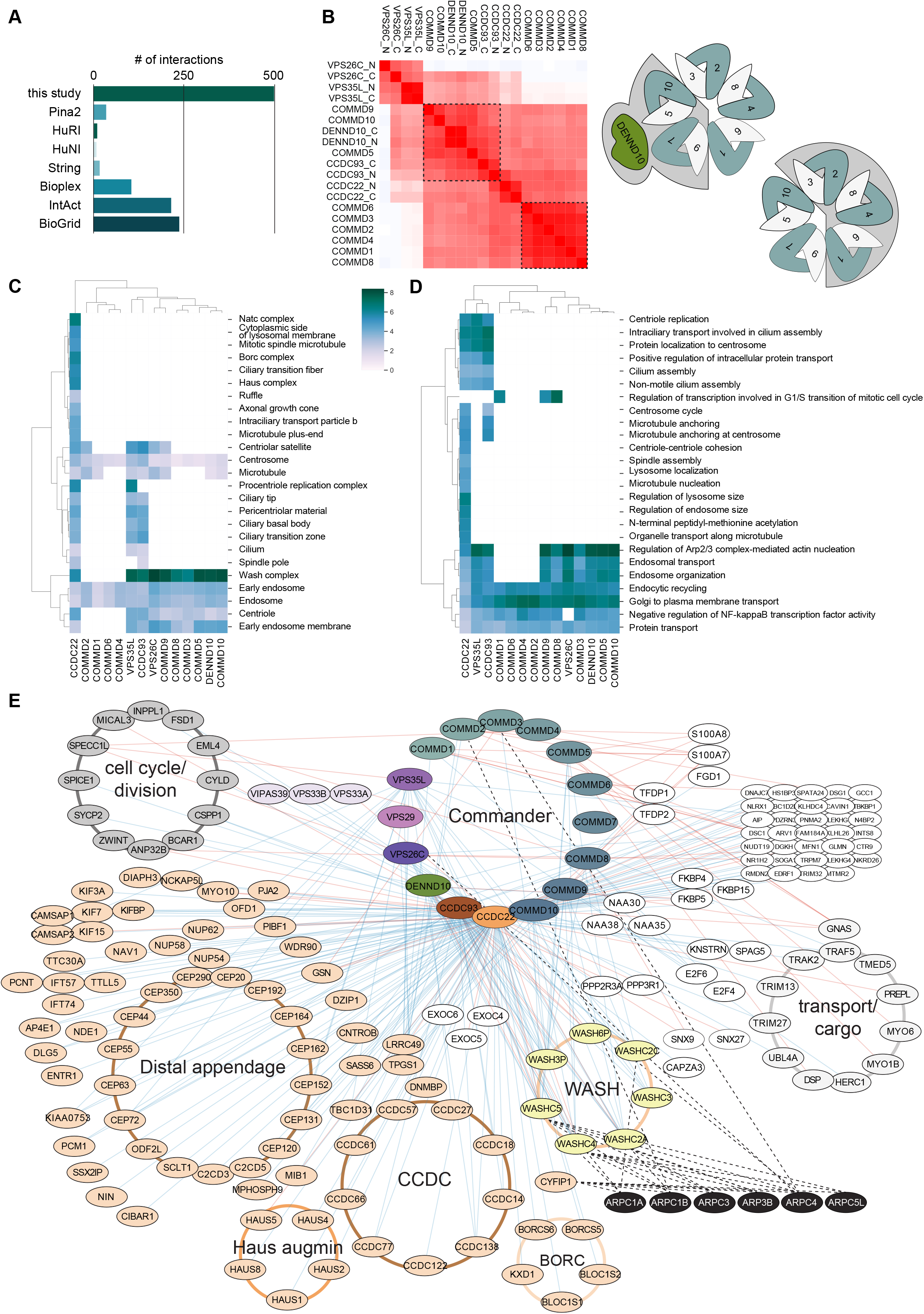
Molecular context, cellular interactions and functions of the Commander complex. (**A**) A comparison of our identified high-confidence Commander complex interactions, identified with the BioID-MS, to reported interactions in the databases. (**B**) Bait-bait clustering of BioID interactions of the Commander proteins interactions reveals their two distinct clusters suggesting two different surface areas, confirmed by the structure. Gene Ontology Cellular Component (**C**) and Biological Process (**D**) term analysis and clustering based on the complex components BioID interactions. (**E**) The complete map of the molecular interactions formed by the Commander complex. Key: The Commander complex components used as the baits are colour coded as in the Figure 1, the identified interactions with AP-MS and BioID are shown as red and blue edges, respectively. The reciprocal interactome analysis with the ARP proteins are shown in the lower right corner. The interacting proteins are organised to designated protein complexes (CORUM; with brown to light orange colored circles), and based on their functions (light grey circles). The nodes linked to cilium assembly (based on GO-BP) are shown with light orange node color.

Subsequently, we conducted bait-bait clustering of BioID interactions of the Commander proteins (**Fig. 5B**), revealing two distinct clusters, suggesting that the Commander complex exhibits two distinct regions. These clusters are similar to the clusters identified by AP-MS, but follow a neat division along WB interfaces. This finding is supported by the complex structure mapped in this study (**Fig. 5B**, upper and lower insets) and indicates that the Commander complex can interact with distinct sets of binding partners on each surface. Moreover, the BioID data can provide information on which side of the Commander complex the other interacting complexes would reside.

We next performed Gene Ontology Cellular Component (GO-CC) and Biological Process (GO-BP) term analysis and clustering based on the complex components’ BioID interactions (**Fig. 5C**). The GO-CC term ‘WASH complex’ was highly (>6-fold) enriched with several of the complex components, but interestingly not detected with COMMD1, 2, 4, and 6, which are on the opposite side of the complex. The terms ‘Early endosome’, ‘Endosome’, and ‘Centrosome’ were moderately enriched for all of the analyzed complex components. We also found that ‘Centriole replication’, ‘Intraciliary transport involved in cilium assembly’, ‘Protein localization to centrosome’, ‘Positive regulation of intracellular protein transport’, ‘Cilium assembly’, and ‘Non-motile cilium assembly’ were significantly enriched for CCDC22, VPS35L, and CCDC93. The CCDC22 also had higher than 4-fold enrichment for several cilia and microtubule associated terms (‘Ciliary tip’, ‘Pericentriolar material’, ‘Ciliary basal body’, ‘Ciliary transition zone’, ‘Cilium, Ciliary transition fiber’, ‘Intraciliary transport particle b’, and ‘Microtubule plus-end’. Additionally the CCDC22 showed an over 6-fold enrichment for the terms of other important complexes (‘Procentriole replication complex’, ‘Wash complex’, ‘Natc complex’, ‘Borc complex’, ‘Haus complex’; **Fig. 5C**).

Additionally, we found that the Commander complex is involved in diverse cellular biological processes (GO-BP; **Fig. 5D**), including ‘Golgi to plasma membrane transport’ (in 100% of components, fold enrichment 3.8–7.4×), ‘Endocytic recycling’ (100%, 3.4–6.2×), ‘Protein transport’ (100%, 2.4–4.5×), ‘Negative regulation of nf-kappab transcription factor activity’ (93%, 2.6–5.8×), ‘Regulation of arp2/3 complex-mediated actin nucleation’ (79%, 5.4–8.4×), ‘Endosomal transport’ (79%, 3.9–6.5×), and ‘Endosome organization’ (79%, 3.8– 6.9×; **Fig. 5D**). All of these are in agreement with the earlier proposed roles of the Commander complex. The fact that COMMD1, 2, 4 and 6 lack these terms is most likely due to their location away from the side of the complex where these interactions take place. Interestingly, ‘Centriole replication’, ‘Intraciliary transport involved in cilium assembly’, ‘Protein localization to centrosome’, ‘Positive regulation of intracellular protein transport’, ‘Cilium assembly’, and ‘Non-motile cilium assembl’ were significantly enriched for CCDC22, VPS35L and for CCDC93. Additionally CCDC22 had enrichment (>3-fold) of a large number of GO-BP terms involved in processes related to centrosomes, microtubules, and intracellular transport and to processes related to membrane-bound organelles (**Fig. 5D, Supplementary Data Table S2**). Although CCDC22 is an integral part of the Commander complex, we cannot rule out the possibility that some of these functions are Commander independent.

### Comprehensive map of the physical and functional interactions formed by the Commander complex

To investigate the molecular interactions formed by the Commander complex in greater detail, we created a comprehensive map that integrated existing protein complex information (from CORUM), pathway information (Reactome Pathway database)(**Fig.S6**), and biological processes (from Gene Ontology) for the Commander complex interactions identified using both AP-MS (red edges/lines) and BioID-MS (blue edges/lines; **Fig. 5E**). The analysis revealed several known complexes that were overrepresented in our data, including the ‘Distal appendage’, ‘HAUS augmin-like complex’, and ‘BORC complex’. Additionally, we identified several other clusters, such as the CCDC protein cluster, cell cycle/division related proteins, and transport/cargo (or cargo transport) proteins. Predicted interactions with the WASH complex were particularly prominent with many of the Commander complex proteins. To further explore this interaction, we re-examined the reciprocal analyses we have performed earlier for the ARPC1A-B, ARPC3, ARP3B, ARPC4, and ARPC5L ^52^. With these baits, we could detect several of the WASH complex components from the opposite direction, as well as a previously unreported interaction of ARPC1B with COMMD2. Overall, these results provide a comprehensive overview of the molecular context, cellular interactions, and functions of the Commander complex, revealing new insights into the complex interactions, organization and functions.

## Discussion

The Commander complex is a highly conserved ubiquitously expressed protein complex that has been implicated in various cellular processes, including protein trafficking, cellular signalling, cell homeostasis, cell cycle, and immune responses. Despite its importance, however, its structure and functions have remained poorly understood. Here, we elucidated the structure and identified the interactions of the endogenous human Commander complex through a combination of cryo-EM and MS-based proteomics. Our results provide the first detailed characterization of the intact, endogenous complex, and a deep characterization of its interactome, 337 novel interactions corresponding to 152 novel interactor in previously unidentified processes such as ‘Cilium Assembly’, Anchoring of the basal body to the plasma membrane’, ‘Organelle biogenesis and maintenance’, ‘Cell Cycle’, M Phase’, and ‘Mitotic Prometaphase’.

Our integrative analyses showed that the Commander complex is organized into two distinct halves, each with its own set of functions. The bottom half of the complex contains two main effectors, that associate the complex to SNX coated PI(3)P-rich membranes and provide a spatiotemporal connection to DENND10 and the WASH complex. The top half of the complex, on the other hand, may act as a cargo recognition site, or as a platform for the assembly of protein complexes in the cytosol via the many binding sites on COMMD NTDs. This structural organization provides the basis for the diverse interactions and roles of Commander complex in cellular processes (**Fig. 6**).

**Figure 6.**
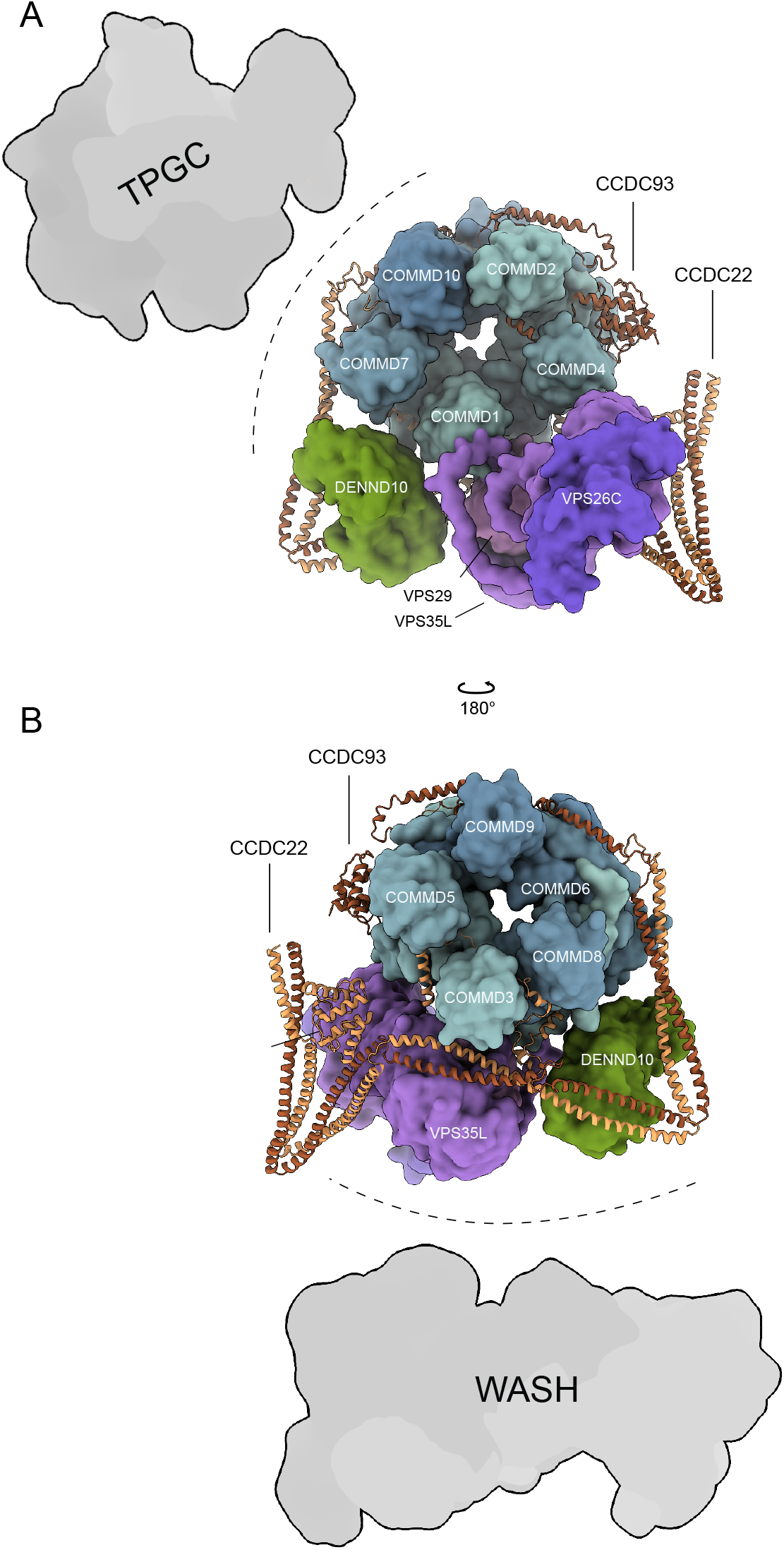
Putative interaction sites of the Commander complex with other complexes. (**A**) Composite model of the Commander complex, indicating putative interaction interfaces with tubulin polyglutamylase complex (TPGC). (**B**) Rotated view of the model in (**A**), with putative interaction interface of the WASH complex indicated.

### Commander interactions with the WASH complex and smaller subcomplexes

Our BioID-MS analysis of the Commander complex identified 148 interacting proteins, the majority of which were previously unreported in six public interactome databases. We found that the WASH complex was highly enriched with several of the Commander complex components, in line with previous reports connecting the Commander and WASH complexes ^78^. We detected the WASH recruitment regulators MTMR2 and PCM1 with similar amounts as the Commander complex components in AP-MS. In contrast, WASH complex components were detected with lower amounts, suggesting a biphasic recruitment of WASH by the Commander complex.

The stable interactions identified with AP-MS in the context of the Commander complex provide valuable insights into the protein-protein interactions and overall architecture of the complex. However, it is important to consider the possibility that the identified interactions may also exist outside of the Commander complex, with complex subunits potentially acting on their own or participating in other smaller subcomplexes or in different complexes altogether. The immediate highlight is the abundant interactions from CCDC22 as the sole Commander component (in both AP-MS and BioID). CCDC22 is also the single subunit implicated in many GO terms and biological processes. Structurally, the termini of CCDC22 and CCDC93 are so close that proximity labelling should overlap significantly. The same principle applies to COMMD2 and COMMD5. Furthermore, AP-MS from the CCDCs should also overlap significantly due to their extended interface. The simplest explanation to this observation is that CCDC22 participates in complexes other than Commander.

### Putative WASH and tubulin polyglutamylase complex binding interfaces

Combined with the structure, our BioID data allows identification of interaction surfaces on the complex. As an example, the most prominent interaction partner in our data is the WASH complex. The majority of BioID signals for WASH comes from CCDC22, CCDC93, DENND10, VPS26C and VPS35L. This indicates that the binding interface is located between DENND10 and Retriever sides of Commander (**Fig. 6B**). This is in line with previous reports showing that WASH complex interacts with Retromer via VPS35 ^79^. This result highlights the ability of systematic proximity labelling to identify interaction surfaces in medium to high molecular weight complexes. Another highlight is an interface consisting of COMMD1, COMMD8, COMMD9, COMMD10 and DENND10. These proteins exhibit BioID signals for two subunits of the tubulin polyglutamylase complex (TPGS1 and LRRC49). In conjunction with COMMD7 (which was not assayed), these proteins come together to form an interaction surface around the I-coil region of the Commander complex (**Fig. 6A**). The identification and characterization of this surface could hold significant implications for understanding tubulin polyglutamylation related cellular processes.

### Possible platforms for cargo-sensing

Structurally, the Commander complex can be divided into two halves: the top half is rigid at the core, while containing some flexible components (NTDs of COMMDs 1, 7, 9 and 10, HLH-motif of CCDC93), and the bottom half which appears in general more heterogeneous in both composition and conformation. The compositional heterogeneity is especially apparent in the Retriever subcomplex which is only completely present in a minor subpopulation of particles in our data. This may be an indication of breakdown of the complex during sample preparation or it may reflect a physiological function. Indeed, COMMD proteins and the CCDCs have been proposed to form a complex without Retriever, termed the CCC-complex ^78^. Interestingly, assuming a similar head-to-head dimerization mode as Retromer, the structure of Commander permits binding of the CCC-complex on a Retriever dimer without steric clashes (**Fig. S5F-H**).

COMMD proteins have been proposed to have a cargo-sensing role in the Commander complex ^19^. This is consistent with the fact that the NTDs of COMMDs are much less conserved than the COMMD domain. In the structural context, we find that the PB-site is a good candidate for interactions with linear peptide epitopes, as several – but not all – of these sites are occupied by such peptides from the CCDCs. The flexibility of the NTDs correlates with the occupancy of this binding site as empty sites on COMMD1 and COMMD9 are the most flexible, followed closely by COMMD7 and COMMD10, each binding a short helix between long flexible regions of CCDC22. At least COMMD9 has been shown to interact with the Notch receptor, a result compatible with the hypothesis that COMMD1 and COMMD9 (and perhaps COMMD7 and COMMD10) could recognize cargo via linear peptide epitopes binding to the PB-site ^19^. Alternatively, these binding sites could act as a platform for accessory complexes.

### Regulation of DENND10 and other Commander components

The role of DENND10 in the complex is not known. In the present structure, it is stably bound on the R-coil, but connection to the I-coil is unclear. Although the binding site identified in the homologous DENND1B with Rab35 is blocked in our consensus structure, the flexibility exhibited by the C-terminal half of DENND10 may expose this binding site. As COMMD1 NTD seems to move in concert with the DENND10, it is tempting to speculate that COMMD1 may regulate the availability of this site through allostery mediated by interactions with cargo. Another possible regulation comes from the phosphoregulation via Ser286 on DENND10 (**Fig. S5A**).

In addition to the phosphorylation site identified for DENND10, we identified various other phosphorylation sites (**Fig. S5A**). Many of these sites coincide with flexible regions in our model (Thr234, Ser301, Ser 305 of CCDC93, Ser70, Ser71, Thr76, Ser80 of VPS35L) which may indicate binding sites for interactors. The cluster in the disordered intervening N-terminal region of VPS35L is particularly interesting, and may represent a regulatory cluster. Other phosphorylation sites are located in folded parts (Ser286 on DENND10, Ser105 on COMMD4, Ser155 on COMMD10 and Ser54 on CCDC22). Perhaps the clearest site for direct binding regulation is the singular site on CCDC22, which is located on the only external face of the domain.

Our GO-CC and GO-BP term analysis and clustering based on the BioID interactions revealed that the Commander complex is involved in diverse cellular biological processes, including Golgi to plasma membrane transport, endocytic recycling, protein transport, and regulation of actin nucleation, among others. Interestingly, the CCDC22 and VPS35L subunits of the Commander complex were significantly enriched for processes related to centrosomes, microtubules, and intracellular transport, indicating that these subunits may play roles in the regulation of these processes.

## Conclusions

Our study provides a comprehensive characterization of the endogenous human Commander complex, including its structural architecture and cellular interactions. Our findings shed new light on the molecular mechanisms underlying the functions of this complex. We validate its role in regulating intracellular trafficking and cell homeostasis, but we also discover previously uncharacterized associations with cilium assembly as well as centrosome and centriole functions. These findings pave the way for further research into the Commander complex, underlying causes of related diseases, in addition to potential drug discovery efforts targeting its components.

### Limitations of the study

Resolution in the bottom half in the Commander complex cryo-EM map limits interpretation of residue level interactions. In addition, the structure of Retriever in the context of Commander could be different compared to the unbound form. These could be addressed by studying individual structures or by incorporating Commander as a part of an even larger complex in order to stabilise mobile parts in future studies.

Future studies could focus on elucidating the specific mechanisms by which the Commander complex regulates the WASH complex and its role in endosomal trafficking. Additionally, further investigations could explore the roles of the CCDC22 and VPS35L subunits in centrosome-related processes and intracellular transport. The identification of previously unreported interacting proteins by our BioID-MS analysis also opens up new avenues for research into the cellular functions of the Commander complex and the potential implications for disease.

Another limitation of our study is that we used human cells, and it is unclear if the Commander complex functions similarly in other organisms. Further studies in model organisms could provide additional insights into the evolution and conservation of the Commander complex and its functions. Additionally, future studies could explore the potential role of post-translational modifications, such as phosphorylation or ubiquitination, in regulating the functions of the Commander complex.

## Acknowledgements

We thank Salla Keskitalo and Antti Tuhkala, Pasi Laurinmäki, Benita Löflund, Dustin Morado and Karin Walldén for technical assistance. The facilities and expertise of the HiLIFE Proteomics and CryoEM units at the University of Helsinki, a member of Instruct-ERIC Centre Finland, FINStruct, and Biocenter Finland are gratefully acknowledged. The cryo-EM data was collected at the Cryo-EM Swedish National Facility funded by the Knut and Alice Wallenberg, Family Erling Persson and Kempe Foundations, SciLifeLab, Stockholm University and Umeå University. The authors wish to acknowledge CSC – IT Center for Science, Finland, for generous computational resources. This study was supported by grants from the Academy of Finland (314669 to J.T.H. and 288475, 319303 and 336470 to M.V.).

## Author contributions

Conceptualization, J.T.H. and M.V.; Writing – Original Draft, E-P.K. and M.V.; Writing - Review & Editing, all authors; Formal analysis of MS and proteomics data, S.L.; Formal analysis of cryo-EM data E-P.K.; Supervision of cryo-EM investigation, J.T.H.; Supervision of MS and proteomics investigation, M.V.; Visualization, E-P.K. and M.V.; Funding Acquisition, J.T.H. and M.V.

## Declaration of interests

The authors declare no competing interests.

## Inclusion and diversity

We support inclusive, diverse and equitable conduct of research.

## Supplemental Figure Titles and Legends

**Supplemental Figure 1. Affinity purification mass spectrometry analysis of the Commander components, related to Figure 1.** (**A**) Dot-plot visualization (BFDR ≤ 0.05) of the Commander complex proteins’ interactors detected by the AP-MS. Each node corresponds to the abundance of the average spectral count for each prey. (**B**) a focused Dot-plot visualization of the CCDC22 interactions. The Commander complex proteins, proteins involved in the WASH complex recruitment and WASH complex components are shown in bold.

**Supplemental Figure 2. Cryo-EM of native and cross-linked Commander, related to Figure 2.** (**A-B**) Representative cryo-EM micrographs and selected 2D class averages of (**A**) native and (**B**) crosslinked Commander. Particle images were low-pass filtered to 4 Å and show particles picked for the consensus map reconstruction. (**C-G**) Cryo-EM data processing workflow for (**C**) native Commander, (**D**) preliminary processing of crosslinked Commander dataset 1, (**E**) final processing of crosslinked Commander datasets 1 and 2 consensus maps, (**F**) focused map 1 of crosslinked Commander datasets (coiled coil region), and (**G**) focused map 2 of crosslinked Commander datasets (Retriever subcomplex).

**Supplemental Figure 3. Molecular models of Commander complex top half, related to Figure 3.** (**A**) AF2 prediction and the predicted alignment error (PAE) plot of the top half of the Commander complex, constituting the full sequences of COMMDs and residues 120–392 of CCDC22 and residues 21–377 of CCDC93. (**B**) *Top*: Example wristbump interface between COMMD5 and COMMD7. *Middle*: three closeups of the model in cryo-EM density, highlighting the residues involved in the wristbump interaction interface between COMMD5 and COMMD7. *Bottom*: schematic representation of the example wristbump interface. Coloring is by sequence conservation within the human COMMD proteins in Top and Bottom subpanels. (**C-D**) Structural models of all (**C**) handshake and (**D**) wristbump interactions. (**E**) Models of NTDs of COMMD proteins (except COMMD6) depicted alongside parts of CCDC93 or CCDC22 that interact with them at the peptide binding site. (**F-G**) Detail of (**F**) CCDC22 α8 or (**G**) CCDC22 α14 binding site on the COMMD ring.

**Supplemental Figure 4. Molecular models of Commander complex bottom half, related to Figure 4.** (**A**) AF2 model of DENND10 region of the Commander complex used as an initial model with predicted alignment error plots indicating the relevant chains. Model is colored according to per-residue pLDDt scores. Model has been trimmed based on the fit to the cryo-EM density map. AF2 model contained all chains of the bottom half during prediction. (**B**) V-coil region of Commander as in (**A**), with different random seed in AF2 prediction. (**C**) Retriever subcomplex model as in **A**. (**D**) Chemical cross-links identified by MS in the context of the Commander structure model. (**E**) Comparison of DENND1B-Rab35 complex structure (PDB 3TW8) with DENND10 in the context of Commander. I-coil sterically blocks the putative Rab binding site on DENND10. (**F**) Structure of Retriever in the context of the Commander complex compared to the Fungal retromer structure. *Interface 1*: VPS29-VPS35(L). *Interface 2:* VPS35(L)-VPS26(C). Retriever adopts a contracted conformation compared to retromer and exhibits larger interaction interfaces.

**Supplemental Figure 5. Detected post-translational modifications, local resolution estimates and putative dimerization mode of Commander complex, related to Figures 1, 3 and 4. (A)** Molecular model of Commander complex with all detected phosphorylation and histidine methylation sites. m: met-His site, p: phosphorylation site. Inset: rotated model showing details on the CCDC93 NN-CH domain side of the complex. (B-E) Local resolution estimates of cryo-EM maps from (B) native Commander, (C) cross-linked Commander consensus map **(D)** cross-linked Commander focused map 1 (Coils) and (E) cross-linked Commander focused map 2 (Retriever). Color bar indicates resolution in Å. (F) Model of putative head-to-head dimerization of Commander complex prepared by superposition via VPS29 and VPS35(L) C-terminal region. (G) Top view of the model in (F). (H) Retromer arch model (PDB ID 6H7W) depicted in the same orientation as Commander dimer model in (F). (I) Top view of the model in (H). Models in (H-I) color-coded as in Fig. S4F.

**Supplemental Figure 6. Molecular interactors, context, and cellular pathways connected with individual Commander complex components, related to Figure 5. (A)** Dot-plot visualization (BFDR ≤ 0.05) of the Commander complex proteins’ interactors detected by the BioID. Each node corresponds to the abundance of the average spectral count for each prey. **(B)** Molecular level localization of the Commander complex proteins obtained by MS-microscopy. **(C)** Reactome pathways enriched for the Commander complex proteins.

## STAR METHODS

## RESOURCE AVAILABILITY

### Lead Contact

Further information and requests for resources and reagents should be directed to and will be fulfilled by the lead contact, Markku Varjosalo.

### Materials Availability

This study did not generate new unique reagents.

### Data and Code Availability

Cryo-EM maps and atomic models have been deposited at Electron Microscopy Data Bank (EMDB) and Protein Data Bank (PDB) and are publicly available as of the date of publication. Accession codes are listed in the key resources table. Collected mass spectrometry data have been deposited at MassIVE.

Any additional information required to reanalyze the data reported in this paper is available from the lead contact upon request. This paper does not report original code.

## EXPERIMENTAL MODEL AND STUDY PARTICIPANT DETAILS

Human: HEK Flp-In T-REx 293 cell line, Invitrogen

The cell line was obtained directly from commercial sources; additionally only low passage cells (passage number <10) were used for experiments. Manufacturers are known to follow the authentication of cells lines batches regularly and certificates of authentication were provided with the cells.

## METHOD DETAILS

### Cloning of Commander complex components

A total of 14 human Commander complex components were obtained from the human ORFeome Libraries (Genome Biology Unit, HiLIFE (University of Helsinki), Horizon Discovery (Perkin-Elmer)). To generate stable isogenic and tetracycline-inducible cell lines, gene constructs were cloned using Gateway® cloning, into N-terminal pTO_HA_StrepIII-N_GW_FRT and N or C-terminal MAC-tagged vectors. After verification by sequencing the constructs were subsequently introduced into Flp-In™ T-REx™ 293 cells (Life Technologies, Carlsbad, CA) as described by Liu et al ^52,53^.

### Cell culture

HEK293 cells have been widely used for the study of protein–protein interactions (PPIs) ^8,54,55^. In this work, their derivative Flp-In™ 293 T-Rex (Invitrogen, Cat# R78007) were used, which allows generating stable cell clones with a single copy of the tagged transgene in their genome ^53^. The cells were cultured in low glucose tetracycline-free DMEM (Sigma Aldrich) supplemented with 10% FBS and 100□μg/ml penicillin/streptomycin (Life Technologies) at 37□°C with 5% CO_2_.

### Affinity purification

For approximately 7□×□10^7^ Flp-In™ T-REx™ 293 cells stably expressing the human Commander complex components, protein expression was induced with 2□μg/ml tetracycline for 24□h (AP-MS and BioID). An additional 50□μM of biotin was added for proximity labelling (BioID). The cells were pelleted using centrifugation, snap frozen in liquid nitrogen, and stored at −80□°C. The samples were then suspended in 3□ml of lysis buffer (50□mM HEPES pH 8.0, 5□mM EDTA, 150□mM NaCl, 50□mM NaF, 0.5% IGEPAL, 1 mM DTT, 1.5□mM Na_3_VO_4_, 1□mM PMSF, 1x protease inhibitor cocktail, Sigma) and lysed on ice for 15 min.

BioID lysis buffer was completed with 0.1% SDS and 80 U/ml benzonase nuclease (Santa Cruz Biotechnology, Dallas, TX), and lysis was followed by three cycles of water bath sonication (3□min) with intervening resting periods (5□min) on ice.

All samples were then cleared by centrifugation, and the supernatants were poured into microspin columns (Bio–Rad, USA) that were preloaded with 200□µl of Strep-Tactin beads (IBA GmbH) and allowed to drain under gravity. The beads were washed 3 times with 1□ml lysis buffer (without SDS for BioID samples) and then 4 times with 1□ml lysis buffer without detergents and inhibitors (wash buffer). The purified proteins were eluted from the beads with 600□µl of wash buffer containing 0.5□mM biotin. To reduce and alkylate the cysteine bonds, the proteins were treated with a final concentration of 5□mM TCEP (tris(2-carboxyethyl) phosphine) and 10□mM iodoacetamide, respectively. Finally, the proteins were digested into tryptic peptides by incubation with 1□µg sequencing grade trypsin (Promega) overnight at 37□°C. The digested peptides were purified using C-18 microspin columns (The Nest Group Inc.) as instructed by the manufacturer. For the mass spectrometry analysis, the vacuum-dried samples were dissolved in buffer A (1% acetonitrile and 0.1% trifluoroacetic acid in MS grade water).

### Mass spectrometry analysis and database searches

The samples were analysed using the Evosep One liquid chromatography system coupled to a hybrid trapped ion mobility quadrupole TOF mass spectrometer (Bruker timsTOF Pro) via a CaptiveSpray nano-electrospray ion source. An 8□cm × 150□µm column with 1.5□µm C18 beads (EV1109, Evosep) was used for peptide separation with the 60 samples per day methods (buffer A: 0.1% formic acid in water; buffer B: 0.1% formic acid in acetonitrile). The MS analysis was performed in the positive-ion mode using data-dependent acquisition (DDA) in PASEF mode with 10 PASEF scans per topN acquisition cycle. Raw data (.d) acquired in PASEF ^56^ mode were processed with MSFragger ^57^ against the human protein database extracted from UniProtKB. Both instrument and label-free quantification parameters were left to default settings.

### Databases to map known interactions

Known interactors were mapped from BioGRID (only experimentally detected interactions) ^58^, Bioplex (interactions with probability over 0.95) ^54^, human cellmap ^59^, IntAct (only experimentally validated physical interactions) ^60^, PINA2 ^61^, and STRING (only with a STRING score□>□0.9) databases ^62^. Domain annotations were mapped from PFam ^63^. Reactome annotations from Uniprot to the lowest pathway level mapping file available at reactome ^64^. Gene ontology and CORUM ^65^ annotations were taken from UniProt. GOCC annotations for CORUM complexes were taken from the CORUM database ^65^.

### Cryogenic electron microscopy (cryo-EM)

For the cryo-EM analyses 2□×□10^9^ Flp-In™ T-REx™ 293 cells stably expressing the N-terminally Strep-tagged human Commander (COMMD9) protein complex were induced with 2□μg/ml tetracycline for 24□h. The cells were pelleted using centrifugation, snap frozen in liquid nitrogen, and stored at −80□°C. The sample was then suspended in 80□mL of lysis buffer (50□mM HEPES pH 8.0, 5□mM EDTA, 150□mM NaCl, 50□mM NaF, 0.5% IGEPAL, 1 mM DTT, 1.5□mM Na_3_VO_4_, 1□mM PMSF, 1x protease inhibitor cocktail (Sigma) on ice. The sample was then cleared by centrifugation, and to remove nucleic acids and intrinsically biotinylated proteins from the sample, 80 U/ml benzonase nuclease (Santa Cruz Biotechnology) and 125 μg/ml of avidin (Thermo-Fisher Scientific) was added to the supernatant followed by second round of centrifugation. The sample was then cleared by centrifugation, and the supernatants were poured into 10 ml gravity-flow columns (Bio–Rad) that were preloaded with 500□µl of Strep-Tactin beads (IBA GmbH) and allowed to drain under gravity. The beads were washed 4 times with 5□ml lysis buffer without protease inhibitors (wash buffer). The purified proteins were eluted from the beads with 3 x 400□µl of low salt buffer (50 mM HEPES pH 8.0, 5 mM EDTA, 40 mM NaCl, 10 mM NaF) containing 0.3□mM biotin and concentrated at +4 °C to 25 μl volume using Amicon ®Ultra 10 kDa MWCO centrifugal filters (Merck Millipore).

To stabilize the Commander complex by cross-linking, 2 mM bis(sulfosuccinimidyl)suberate (BS3; Thermo Scientific) was added for the final 30 min of the concentration. The crosslinked complex was purified in size exclusion chromatography, using Sepharose 6 column in ÄKTA Pure purification system using low salt buffer.

Samples were supplemented with DM at a final concentration of 0.85 mM. Final protein concentration was 0.5 mg/ml (native Commander) or 0.1 mg/ml (cross-linked Commander). A 3-µl aliquot of sample was applied on a glow-discharged Quantifoil Cu 200 mesh R1.2/1.3 grid, followed by incubation period and plunge-freezing in liquid ethane using a VitroBot Mark IV cryoplunger (Thermo Fisher Scientific). Blot time of 7 s at 100% relative humidity and 6°C temperature was used with a blot force of −15. Cross-linked Commander grids were prepared using Quantifoil Cu 200 mesh R1.2/1.3 grids with an additional 2-nm amorphous carbon support film and an incubation time of 5 min on the grid. The grids were stored in liquid nitrogen for subsequent screening and imaging.

Data for the native Commander complex was collected at the Umeå Core Facility for Electron Microscopy, Sweden, using a Titan Krios 300 kV transmission electron microscope (Thermo Fisher Scientific), with an X-FEG electron source and a Gatan K2 direct electron detector camera equipped with a BioQuantum energy filter. Data was collected at 1.5 to 3.0 µm defocus, 40 frames at 42.8 e^−^/Å^2^ total electron exposure and a pixel size of 0.82 Å for a total of 5,884 movies collected. Energy filter slit width was set at 20 eV. For cross-linked Commander, two datasets were collected at the Cryo-EM Swedish Infrastructure Unit of SciLifeLab, Stockholm, Sweden, in two separate sessions from two grids prepared identically in the same session, using a Titan Krios 300 kV transmission electron microscope (Thermo Fisher Scientific), with an X-FEG electron source and a Gatan K3 direct electron detector camera equipped with a BioQuantum energy filter set at 20 eV slit width. Dataset 1 was collected at 0.4 µm to 2.2 µm defocus, 50 frames at 59 e^−^/Å^2^ total electron exposure and a pixel size of 0.846 Å for a total of 20,675 movies collected. Dataset 2 was collected at 0.6 µm to 2.0 µm defocus, 45 frames at 56 e^−^/Å^2^ total dose and a pixel size of 0.862 Å for a total of 35,094 movies collected. Data collection parameters are in **Supplemental Table 1**.

Cryo-EM data were processed in CryoSPARC v.4.1.1 unless stated otherwise ^66^. Native Commander complex was processed in tandem with the crosslinked dataset 1 (see below, **Fig. S2C**). Movies were motion corrected and contrast transfer function (CTF) parameters estimated using Patch Motion Correction, and Patch CTF Estimation in cryoSPARC, respectively ^66^. After manual curation of micrographs, 5748 exposures were retained. Particles were initially picked using template picking, using an initial volume from preliminary analysis carried out with cross-linked dataset 1 as a picking template. A total of ∼867,000 particles were picked and cleaned by four rounds of 2D classification, resulting in ∼127,000 particles retained. This particle set was binned by a factor of 2 and refined by one round of ab initio reconstruction using four classes, followed by heterogeneous refinement against the ab initio volumes and homogeneous refinement of the class with highest resolution features (∼51,000 particles, 6.8 Å resolution, Nyquist-limited). This set was further refined by a similar round of ab initio reconstruction using four classes followed by heterogeneous refinement and homogeneous refinement of the top class (∼17,000 particles, 6.8 Å resolution, Nyquist-limited). This was followed by per-particle motion correction ^67^ using unbinned particles and homogeneous refinement (∼17.000 particles, 3.7 Å resolution). The resulting particles were used to train a TOPAZ model ^68^ and re-pick the micrographs. A total of ∼340,000 particles were extracted, followed by a single round of 2D classification and removal of duplicate picks, which yielded a set of ∼137,000 particles. Heterogeneous refinement against four copies of the final volume from the initial processing yielded a model with ∼90,000 particles and 3.3-Å resolution from non-uniform refinement ^69^. Particles were further unbinned and local motion correction was applied, which resulted in a consensus map at 3.2-Å resolution and an estimated B-factor of 66.

For cross-linked Commander, datasets 1 and 2 were processed separately until the final consensus refinement stage (**Fig. S2D**). Before the final processing workflow, an initial exploratory analysis was carried out on dataset 1 where particles were picked using the supervised picking routine of xmipp3 from micrographs denoised using the Noise2Noise implementation in crYOLO ^70^ within the Scipion framework ^71^. These particles were then processed in cryoSPARC by iterative cycles of multi-class ab initio and heterogeneous refinement to yield an initial model that encompassed the maximum volume of the complex at the cost of resolution (6.6 Å, ∼24,000 particles). The final particles were used to train a TOPAZ neural network model for auto-picking particles in both datasets 1 and 2.

Movies were motion corrected and CTF parameters estimated using patch motion correction and patch CTF estimation implementations in cryoSPARC. Micrographs were manually curated to remove bad micrographs with a resulting micrograph count of 19,630 for dataset 1. The TOPAZ network trained using the initial processing was used to pick ∼2.9 million particles. Particles were extracted in a box of 560×560 pixels at full size, downsampled to 140×140 pixels for 2D classification. Two rounds of 2D classification were carried out to classify the particles into 100 classes, with ∼872,000 particles retained. The resulting particles were re-extracted in a box of 280×280 pixels and used in a 6-class ab initio reconstruction followed by heterogeneous refinement using the 6 ab initio models as template volumes. One class containing the highest resolution features was selected, containing ∼397,000 particles. These particles were used at full size for non-uniform refinement, optimizing per-particle scales, defoci and per-group CTF parameters with tilt and trefoil fits.

Dataset 2 was processed similarly to dataset 1 (**Fig. S2D**). Pixel size set to 0.846 Å / pix to facilitate merging of the data (calibrated pixel size 0.861 Å / pix). After micrograph curation, 33,879 micrographs were retained. TOPAZ picking yielded ∼4.7 million particles, reduced to ∼4.5 million after one round of 2D classification. Heterogeneous refinement using the 6 ab initio volumes from dataset 1 yielded a single class with high resolution features, with ∼1.0 million particles retained. This class was first refined using non-uniform refinement and further polished by a round of 4-class ab initio reconstruction followed by heterogeneous refinement. The final particle stack, containing ∼577,000 particles, was merged with the particles from dataset 1 (total of ∼958,000 particles) and refined using non-uniform refinement with per-particle defocus optimization and per-group CTF parameter optimization, fitting tilt and trefoil parameters and per-particle scales. The combined particles were polished by heterogeneous refinement against the current consensus map and 6 ab initio maps from the 0th iteration against the data used as junk sinks. The particles that ended up in the consensus class (∼667,000) were refined locally with a mask encompassing the COMMD-ring. The final gold-standard global CTF resolution was estimated at 2.9 Å with an estimated B-factor of 77. Local motion correction was performed but this showed no improvement in resolution.

To find the substructure of the bottom half of Commander (focused map 1, **Fig. S2F**), partial signal subtraction was carried out using the final COMMD ring map and a mask encompassing the COMMD ring region of the map, using a low-pass filter of 10 Å with the final particle set. The resulting subtracted particles were refined locally using a gaussian prior for poses and shifts at standard deviation of prior over rotation of 30° and standard deviation of prior over shifts of 14 pix with FSC noise substitution enabled and a maximum alignment resolution of 12 Å imposed and using a batch size of 5000, resulting in a map with nominal resolution of 8.6 Å. This particle set was then polished with several rounds of heterogeneous refinement against six junk sink classes, followed by local refinements, to reach a final particle stack of ∼125,000 particles at a nominal resolution of 6.5 Å.

To resolve the Retriever substructure (focused map 2, **Fig. S2G**), partial signal subtraction was carried out using the final COMMD ring map and a mask encompassing the I-coil, half of R-coil and DENND10 regions of the map using a low-pass filter at 10 Å with the final particle set. The resulting particles were refined locally using a gaussian prior for poses and shifts at standard deviation of prior over rotation of 20° and standard deviation of prior over shifts of 10 pix with FSC noise substitution enabled and a batch size of 5000, resulting in a map with nominal resolution of 7.8 Å. The initial model was the coils map obtained from the previous step. This map was then used as an input in heterogeneous refinement together with six junk sinks as described for the consensus map to remove bad particles, resulting in a particle stack of ∼195,000 particles. This particle set was refined locally using settings as above to reach a resolution of 7.6 Å. 3D classification with 10 classes, 250 Å high-pass filter, O-EM batch size of 2000 per class and PCA initialisation mode with default settings was performed and the class with most features at the VPS26C end of the map was selected. This particle set of ∼13,000 particles was locally refined with an additional 10 Å maximum alignment resolution to prevent overfitting, reaching a nominal resolution of 7.5 Å.

3D variability analysis was implemented in cryoSPARC ^72^. Particles used in reconstruction of focused map 1 were used without subtraction and with 3D-alignments from the consensus map reconstruction. 20 volumes along each principal component were reconstructed using the “intermediates” option.

Atomic models were built into the cross-linked Commander cryo-EM density using AlphaFold2 models as a starting point in COOT ^73^ and adjusted using ISOLDE ^50^. AF2 models were generated using a local installation of ColabFold v. 1.5.2 ^47^ running AlphaFold v.2.3.1 ^48^ and AlphaFold Multimer v3^49^ (Predictions were run using 10 recycles, 1 ensemble, with dropouts activated, max-seq of 128 and max-extra-seq 256 with relaxation disabled. AF2 output models were fitted into density as rigid bodies, and predictions or parts of predictions most closely matching the density were selected for model building. The top half of Commander was predicted using full sequences of COMMD proteins as well as residues 120–392 of CCDC22 and 21–377 of CCDC93. The CCDC sequences included the I-coil region, which was necessary for correct placement of α15 and α16 of CCDC22, although the orientation of the I-coil itself in these predictions showed no correspondence to the cryo-EM density. The initial models for DENND10 and the I, R, and V-coils were obtained from two separate predictions (different random seed) from the same run where the sequences of CCDC22 (residues 1–110, 332–627), CCDC93 (residues 313–631), DENND10, VPS29, VPS35L and VPS26C were used as input. Output models were split such that the DENND10 side of the model (**Fig. S4A**) contained residues 313–425 of CCDC93, residues 322–436 of CCDC22 and DENND10, while the V-coil side of the model (**Fig. S4B**) contained residues 426–631 of CCDC93 and residues 1–110 and 437–627 of CCDC22. The structure of the Retriever subcomplex was predicted separately (**Fig. S4C**). For parts in lower resolution areas of the map (NTDs of COMMD1, COMMD7, COMMD9 and COMMD10, CCDC22 and CCDC93 in the lower half, DENND10 and Retriever) were modelled in ISOLDE with corresponding AF2 input models as distance and torsion restraints, with predicted-alignment error weighting enabled. The ISOLDE-adjusted models were refined in real space in Phenix ^74^ against sharpened maps, using the parameters created in ISOLDE with command “isolde write phenixRsrInput”. Structures were validated using Phenix and MolProbity ^75^. Figures were made using ChimeraX ^76^. For regions with less than 4 Å resolution, poly-alanine models were modelled and sidechain information is presented in figures for visualization purposes only. Model refinement parameters are in **Supplemental Table 2**.

## QUANTIFICATION AND STATISTICAL ANALYSIS

Significance Analysis of INTeractome (SAINT) express version 3.6.0 ^77^ and Contaminant Repository for Affinity Purification (CRAPome, http://www.crapome.org) were used as statistical tools for identification of specific high-confidence interactions from AP-MS and BioID data. 17 control runs with MAC-tagged GFPs were used as controls for SAINT analysis. Identifications with a SAINT-assigned Bayesian FDR ≥□0.05 were dropped, as well as any proteins that were detected in ≥□20% of CRAPome experiments, unless the spectral count fold change was over 3 when compared to CRAPome average. The remaining HCIs were then used for further analysis.

### Supplemental Data Tables

Supplemental Data Table 1. High-confidence interactions detected via AP-MS and BioID for the 14 Commander complex components.

Supplemental Data Table 2. Functional classifications of the Commander complex interactors, and detected phosposites and cross-links for the Commander complex proteins.

### Supplemental Movies

Supplemental Movie 1. Atomic model of the Commander complex. Related to figure 6.

## Notes

### Competing Interest Statement

The authors have declared no competing interest.

