## Supplementary figures and images for "Structure and Interactions of the Endogenous Human Commander Complex"

### Supplemental Figure 1

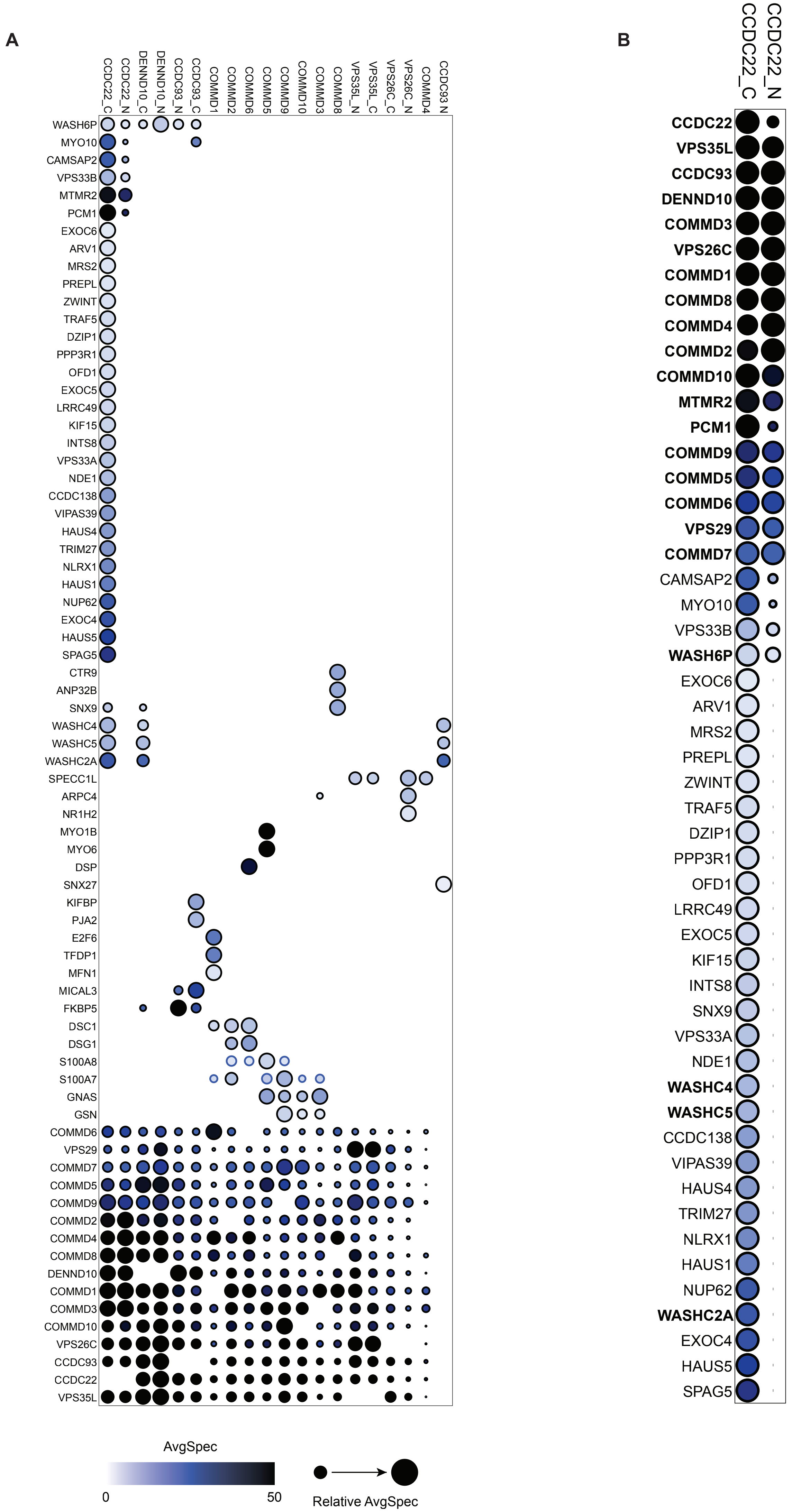

### Supplemental Figure 2

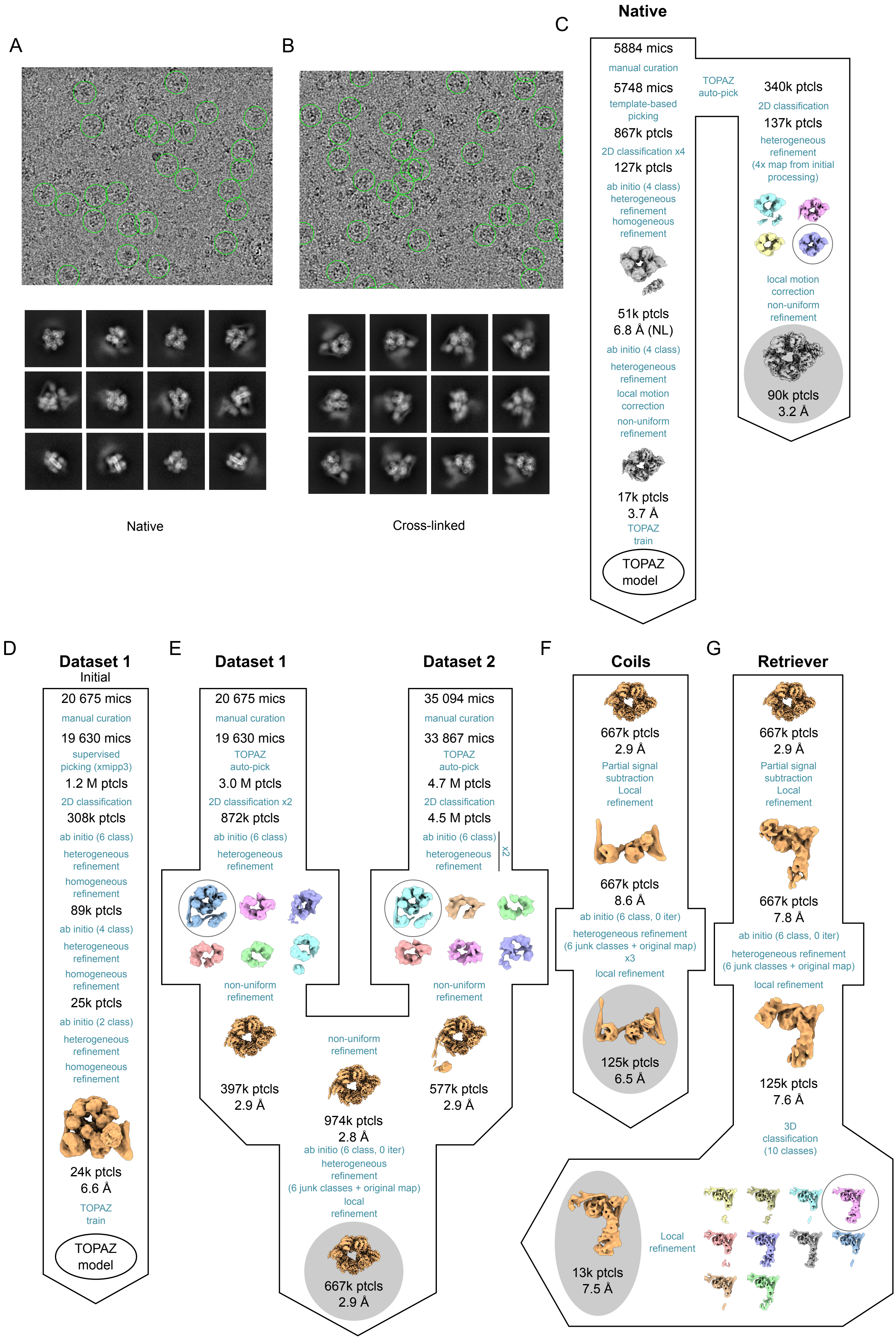

### Supplemental Figure 3

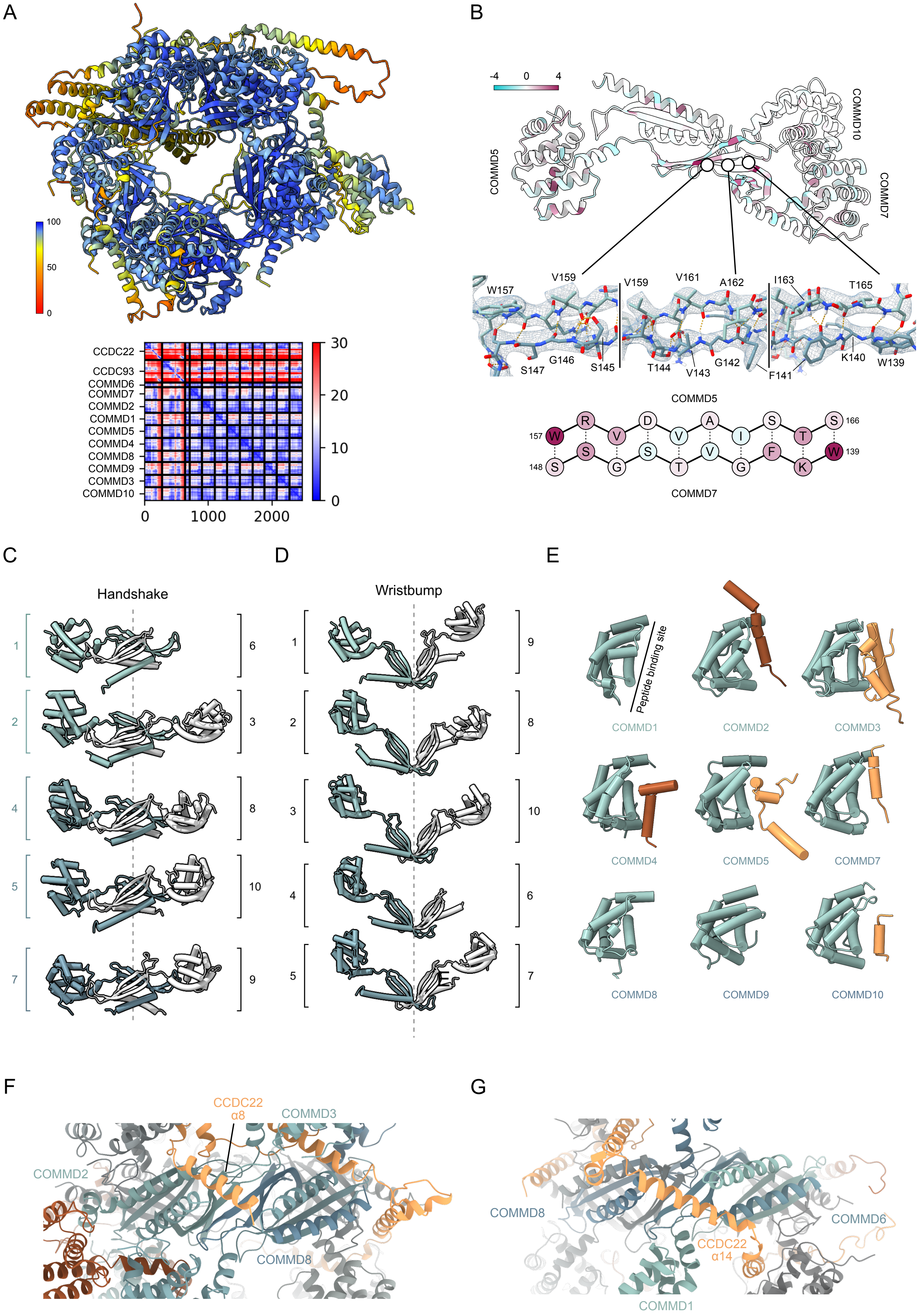

### Supplemental Figure 4

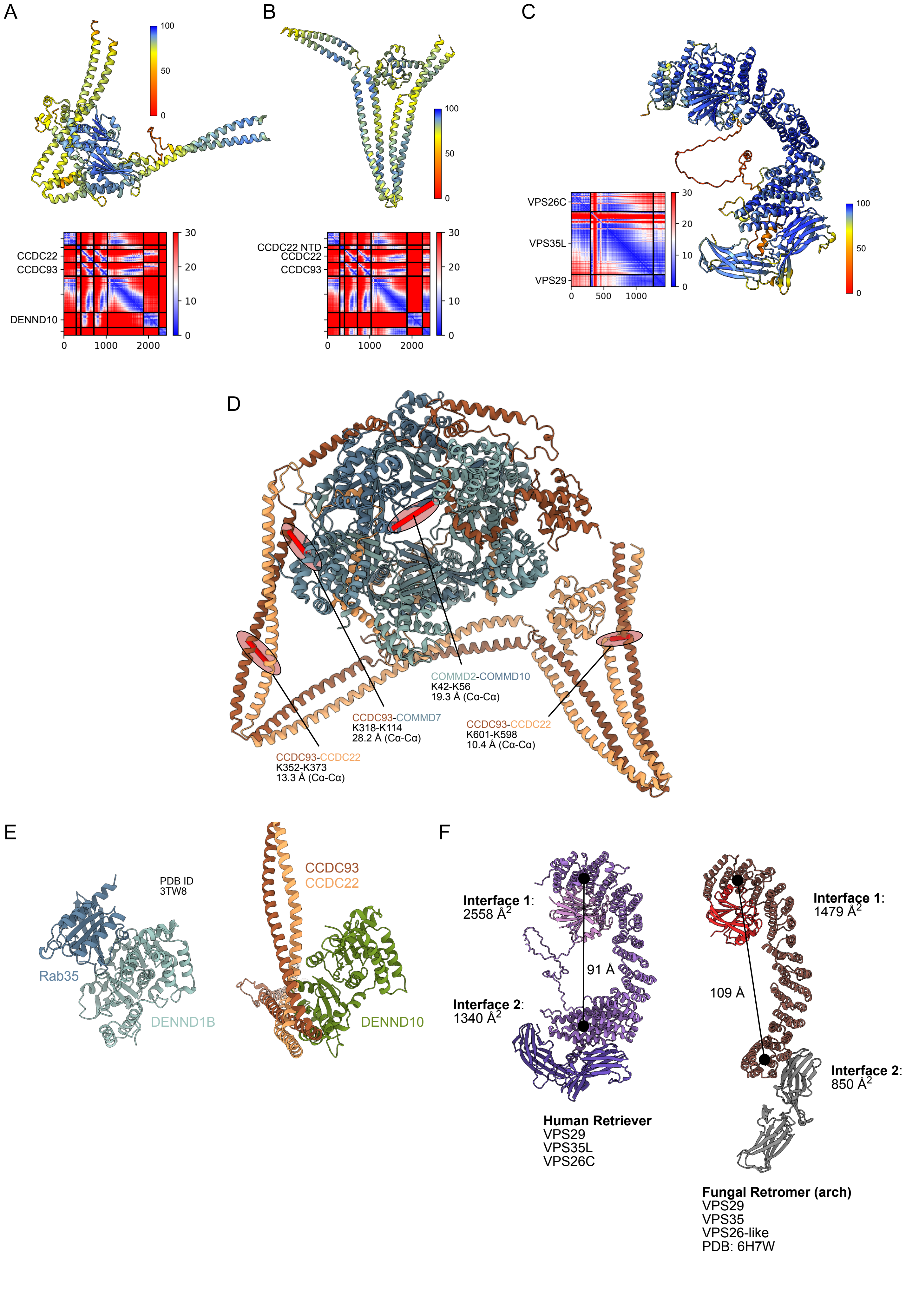

### Supplemental Figure 5

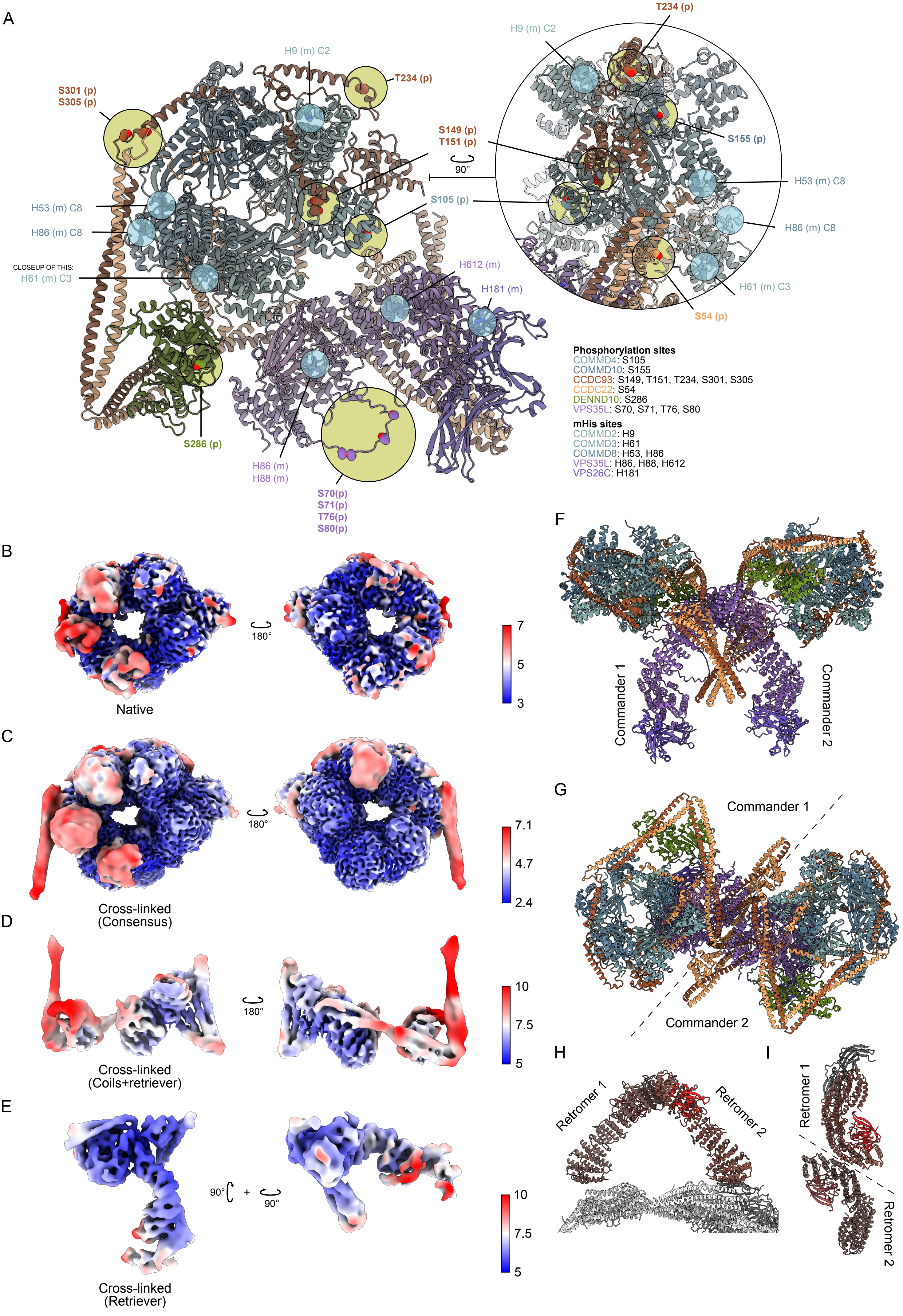

### Supplemental Figure 6

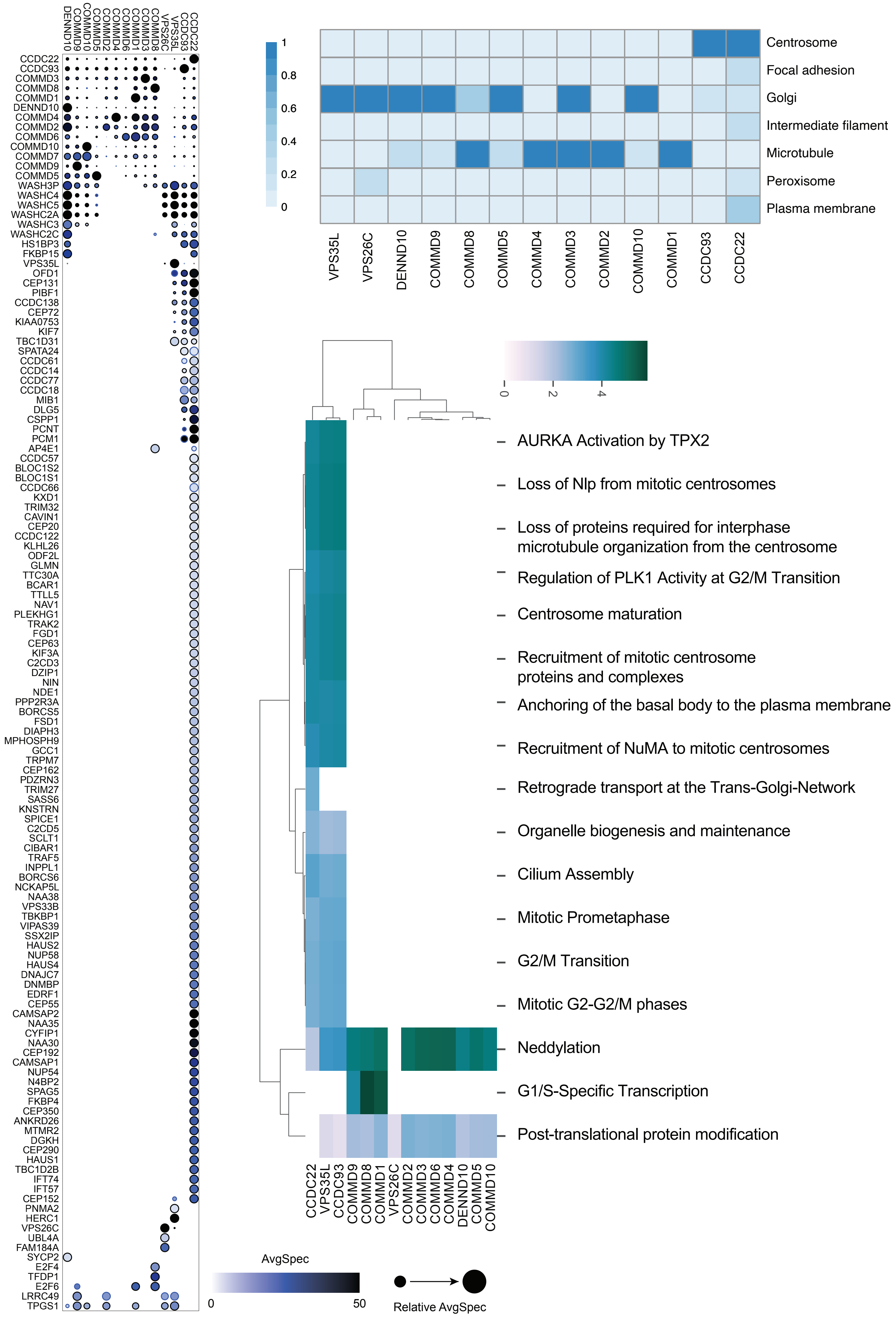
